# Hypothalamic GABRA5-positive Neurons Control Obesity via Astrocytic GABA

**DOI:** 10.1101/2021.11.07.467613

**Authors:** Moonsun Sa, Eun-Seon Yoo, Wuhyun Koh, Mingu Gordon Park, Hyun-Jun Jang, Yong Ryoul Yang, Jiwoon Lim, Woojin Won, Jea Kwon, Mridula Bhalla, Heeyoung An, Yejin Seong, Seung Eun Lee, Ki Duk Park, Pann-Ghill Suh, Jong-Woo Sohn, C. Justin Lee

**Author notes:** Correspondence: C. Justin Lee.

## Abstract

The lateral hypothalamic area (LHA) regulates food intake and energy expenditure. Although LHA neurons innervate adipose tissues, the identity of neurons that regulate fat is undefined. Here we identify that Gabra5-positive neurons in LHA (GABRA5^LHA^) polysynaptically project to brown and white adipose tissues in the periphery. GABRA5^LHA^ are a distinct subpopulation of GABAergic neurons and show decreased pacemaker firing in diet-induced obesity (DIO) mouse model. Chemogenetic inhibition of GABRA5^LHA^ suppresses energy expenditure and increases weight gain, whereas gene-silencing of Gabra5 in LHA decreases weight gain. In DIO mouse model, GABRA5^LHA^ are tonically inhibited by nearby reactive astrocytes releasing GABA, which is synthesized by MAOB. Gene-silencing of astrocytic MAOB in LHA reduces weight gain significantly without affecting food intake, which is recapitulated by administration of a MAOB inhibitor, KDS2010. We propose that firing of GABRA5^LHA^ facilitates energy expenditure and selective inhibition of astrocytic GABA is a molecular target for treating obesity.

## INTRODUCTION

Obese people have an imbalance in food intake and energy expenditure which are regulated by neural circuits which work inside the hypothalamus and extend beyond the hypothalamus (Kong et al., 2012; Thaler et al., 2012). Hypothalamus consists of a number of small nuclei which include LHA. Although LHA occupies an extended field of the hypothalamus, it is substantially less anatomically defined (Bernardis and Bellinger, 1993; Flament-Durand, 1980; Palkovits et al., 1980; Stuber and Wise, 2016). A subpopulation of LHA neurons are known to innervate brown adipose tissue (BAT) and white adipose tissue (WAT) to mediate thermogenesis in BAT, browning of WAT and energy storage in WAT (Cerri and Morrison, 2005; Contreras et al., 2017). However, the precise cell types that innervate BAT and WAT to mediate thermogenesis and energy storage are still under active investigation. The LHA contains several cell types expressing different transmitters and hormones, including neurons expressing melanin-concentrating hormone (MCH) and hypocretin/orexin (hcrt/orx) (Bittencourt, 2011; Lee et al., 2021; Sakurai et al., 1998); MCH neurons in LHA negatively regulate BAT activity to suppress energy expenditure (Oldfield et al., 2002), whereas orexin neurons send excitatory projections to increase BAT activity and energy expenditure with decreasing in food intake (Berthoud et al., 2005; Contreras et al., 2015; Kakizaki et al., 2019; Tupone et al., 2011; Zink et al., 2018). In addition, LHA contains other neurons that express neither MCH nor orexin (Backberg et al., 2004; Karnani et al., 2013; Kosse et al., 2017): A large population of GABAergic neurons in LHA are intrinsically depolarized and distinct from MCH and orexin (Karnani *et al*., 2013). These GABAergic neurons are defined by the presence of components necessary for GABA synthesis and release, including GAD65, GAD67 and vesicular GABA transporter (VGAT) (Bonnavion et al., 2016; Hassani et al., 2010; Jennings et al., 2013; Shin et al., 2007). Due to their location in LHA, these neurons might function as critical regulators of energy balance. However, the further classification and functional characterization of this vast majority of GABAergic neurons in LHA are still unexplored (de Vrind et al., 2019). Furthermore, how this heterogeneous population of GABAergic neurons in LHA interact with other cell types remains poorly understood.

It has been reported that extracellular GABA level in mediobasal of hypothalamus becomes elevated during chronic over-nutrition (Zhang et al., 2017). However, how this extracellular GABA is synthesized and what causal role it plays in the pathogenesis of obesity and related metabolic syndrome remains to be elucidated. We have previously shown that monoamine oxidase B (MAOB), mainly expressed in astrocytes, synthesizes astrocytic GABA (Chun et al., 2018; Chun and Lee, 2018; Yoon et al., 2014). MAOB mediates degradation of polyamine putrescine, which is a byproduct of toxin degradation, to generate GABA in astrocytes (Yoon *et al*., 2014). Notably, MAOB is elevated in transcriptionally profiled LHA cells in high fat diet (HFD)-fed mice in recent study (Rossi et al., 2019). Astrocytes are known to be actively involved in the regulatory aspects of metabolic control, such as feeding and brain glucose uptake (Bouyakdan et al., 2019; Chari et al., 2011; Chen et al., 2016; Garcia-Caceres et al., 2016; Kim et al., 2014b; McDougal et al., 2013; Varela et al., 2021; Yang et al., 2015). In addition to physiological condition, increasing lines of evidence point to an involvement of hypothalamic astrocytes in the pathogenesis of DIO (Gonzalez-Garcia and Garcia-Caceres, 2021). Reactive astrocytes are observed in several regions of hypothalamus after HFD feeding (Buckman et al., 2013). Consumption of dietary fats also induce metabolic damages on hypothalamic neurons, such as neuronal injury and a reduction of synaptic inputs in LHA (Lizarbe et al., 2019; Moraes et al., 2009; Thaler *et al*., 2012). However, how these neuronal dysfunctions are mediated by GABA from reactive astrocytes in LHA remains elusive. Furthermore, the role of reactive astrocytes in LHA as the controller of pathological processes by over-nutrition still remains undetermined.

The released GABA from reactive astrocytes, as conventionally called tonic GABA, is mediated by extrasynaptic GABA_A_ receptors in neighboring neurons (Yoon et al., 2012). Tonic GABA reduces spike probability of granule cells in Alzheimer’s disease (AD) mouse models (Jo et al., 2014). Extrasynaptic GABA_A_Rs composed of α5βγ2, α4βδ, α6βδ and α1βδ subunits and are located largely at extrasynaptic sites (Brickley and Mody, 2012; Caraiscos et al., 2004). It has been previously demonstrated that α5 subunit of GABA_A_R (Gabra5) shows moderate expression, whereas δ subunit shows faint expression in LHA (Hortnagl et al., 2013). It has been further demonstrated that Gabra5-positive neurons are distinct from MCH, orexin A and orexin B in LHA (Thaler *et al*., 2012). However, whether these Gabra5-positive neurons in LHA (GABRA5^LHA^) are GABAergic and their physiological role in energy expenditure and food intake are entirely unknown.

In this study, we hypothesized that GABRA5^LHA^ have regulatory role in energy balance by interacting with astrocytes via gliotransmitter GABA. Based on genetic, pharmacological and electrophysiological approaches, we found that GABRA5^LHA^ are a distinct sub-population of pacemaker firing GABAergic neurons, regulating energy expenditure without affecting food intake via astrocytic GABA in HFD-fed mice. Given these findings, we suggest that genetic and pharmacological inhibition of excessive astrocytic GABA synthesis and GABA_A_ receptor containing α5 subunit could be effective therapeutic strategies for obesity.

## RESULTS

### Pacemaker firing in GABAergic GABRA5^LHA^ is decreased in HFD mice

To identify and characterize functional roles of GABRA5^LHA^, we developed Gabra5 promoter-containing virus and injected AAV-mGabra5-EGFP-Cre into the LHA (Figures 1A and 1B). Next, we performed immunohistochemical staining with antibodies against Gabra5, orexin A, orexin B, MCH and GABA in LHA-injected slices to confirm the specificity of the promoter. Most of Gabra5 promoter-containing EGFP-positive cells were overlapped with Gabra5 and GABA, but not with orexin or MCH (Figures 1C and 1D). These results suggest that Gabra5 promoter specifically target GABRA5^LHA^ which are mostly GABA-producing neurons.

**Figure 1.**
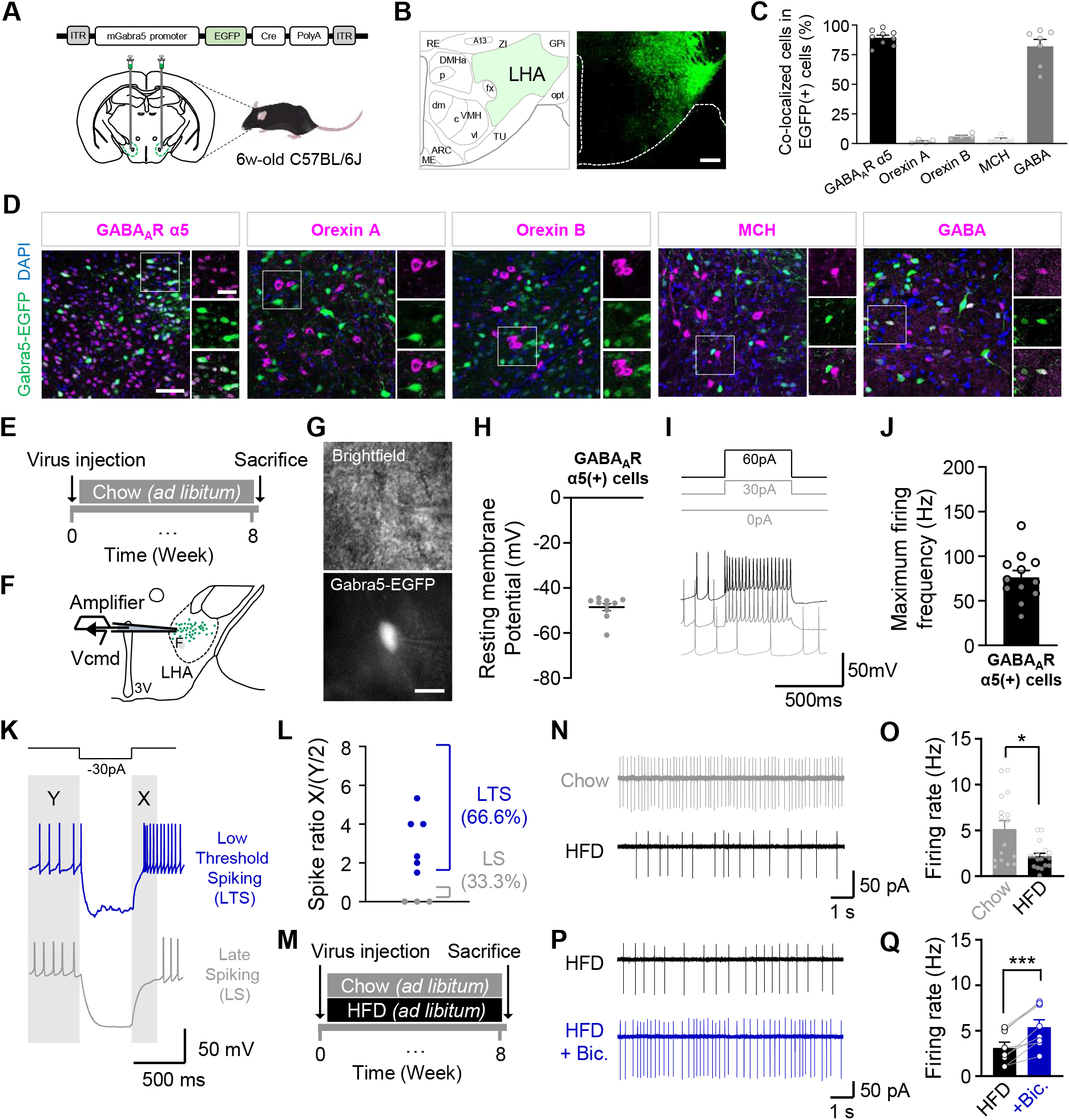
GABAergic GABRA5^LHA^ show decreased activity in HFD. (A and B) Experimental schema and representative image for EGFP neurons infected with AAV-Gabra5-EGFP-cre in LHA (n=4 mice). Bregma, −1.58 mm AP. Scale bar, 100 μm. (C) Quantification of colocalized cell in EGFP neurons. (D) Gabra5-EGFP is colored by green and Gabra5, orexin A, orexin B, MCH and GABA signals are colored by Magenta. Left, representative image with inset of region. Scale bar, 100 μm. Right, magnification of ROI in LHA regions. Scale bar, 50 μm. (E and F) Timeline and schema of whole cell patch clamp recording in GABRA5^LHA^. (G) Above, contrast and bottom, fluorescence micrograph of GABRA5^LHA^ neuron being recorded. Scale bar, 10 μm. (H) Quantified resting membrane potentials(mV) of GABRA5^LHA^ based on I–V. (I) Representative traces of non-fast spiking GABRA5^LHA^ with depolarizing current steps. (J) Maximal firing frequency of GABRA5^LHA^ when a maximally depolarizing current step is applied. (K) Representative response of top, low threshold spiking (LTS) and bottom, late spiking (LS) to the current step. Classification of non-fast spiking cells by hyperpolarizing current steps. (L) Summary of spike ratio calculated from the X (250 ms) and Y (500 ms) time windows as spike amounts during X/(Y/2). (M) Timeline for cell-attached patch-clamp recording in GABRA5^LHA^ experiment. (N) Representative traces of cell-attached recording in GABRA5^LHA^ from chow diet (top) and high fat diet (bottom) mouse. (O) Quantification of firing rate between chow and HFD. n=3 mice per group. (P) Representative traces of cell-attached recording in GABRA5^LHA^ from HFD (top) and HFD with bicuculline (bottom). (Q) Quantification of firing rate between HFD and bicuculline added. Paired t test. Data represents Mean ± SEM. *, p<0.05; **, p<0.01; ***, p<0.001.

To investigate the intrinsic electrical properties of GABRA5^LHA^, we used whole-cell patch-clamp recordings in acute brain slices of AAV-Gabra5-EGFP-Cre injected mice (Figures 1E-1G). GABRA5^LHA^ were spontaneously active in slice with near −50mV of resting membrane potential (Figure 1H). We measured action potential (AP) wave forms for GFP-positive neurons, using sustained current injection (Figure 1I). GABRA5^LHA^ were spiking at near-threshold membrane potential and showed maximum firing frequency of ∼76 Hz during maximal depolarization (Figure 1J). It has been previously shown that LHA GAD65 cells fall into four subtypes based on their distinct electrical signatures with evoked firing: fast-spiking (FS), late-spiking (LS), low-threshold spiking (LTS) and regular-spiking (RS) (Karnani *et al*., 2013). Based on this categorization, we characterized the firing patterns of GABRA5^LHA^ before and after a hyperpolarizing current step (Figure 1K). Approximately 66% of GABRA5^LHA^ showed increased firing frequency after the step by more than 50% of that before the step. About 33% of GABRA5^LHA^ did not start firing immediately after the step, but rather showed a slow ramp depolarization, resulting in a firing rate of less than 50% before the step (Figures 1K and 1L). We found that GABRA5^LHA^ fell into two major subtypes based on evoked firing: LS and LTS (Figure 1L). These results indicate that GABRA5^LHA^ are LS- and LTS-subtypes of GAD65-positive GABAergic neurons in LHA.

To investigate the diet-induced changes in GABRA5^LHA^, we performed loose cell-attached patch clamping on GABRA5^LHA^ without disturbing the intracellular ionic concentration in acutely prepared LHA slices from 8-week HFD-fed mice (Figure 1M). We found that the GABRA5^LHA^ pacemaker firing rate was significantly decreased in HFD mice (Figures 1N and 1O). The decreased firing rate was significantly restored by treatment with GABA_A_ receptor antagonist, bicuculline (50 μM), in HFD mice (Figures 1P and 1Q). These results indicate that GABA-mediated inhibition tonically suppresses the GABRA5^LHA^ pacemaker firing in HFD mice.

### Chemogenetic inhibition of GABRA5^LHA^ suppresses energy expenditure

We next investigated whether neuromodulation of GABRA5^LHA^ regulates body weight and food intake by using chemogenetics to inhibit these neurons. We used the designer receptor exclusively inhibited by designer drug (DREADD) hM4Di with a combination of AAV-mGabra5-EGFP-cre and AAV-hSyn-DIO-hM4Di-mCherry viruses (Figures 2C and S1A-S1D). After virus injection in LHA (Figure 2A), the mice were fed with HFD and administered with clozapine N-oxide (CNO) by drinking for 5 weeks (Figure 2B). Inhibition of GABRA5^LHA^ by drinking CNO led to a significant increase in body weight and food intake starting from 4 weeks (Figures 2D-2F). Then we placed the mice in metabolic cages to enable automated phenotyping of whole animal metabolic activity using the comprehensive lab animal monitoring system (CLAMS). Inhibition of GABRA5^LHA^ led to a significant decrease in total energy expenditure (Figures 2G and 2H), carbon dioxide production (Figures 2I and 2J) and oxygen consumption (Figures 2K and 2L) in dark cycle without changes in locomotor activity (Figures 2M and 2N). These results imply that GABRA5^LHA^ facilitate energy expenditure in HFD mice. Then, we examined the expression of candidate genes which are related to thermogenesis in adipose tissues by qRT-PCR analysis using reference gene as 18S ribosomal RNA (Cannon and Nedergaard, 2004; Cero et al., 2021; Contreras *et al*., 2015; Kurylowicz et al., 2015; Orozco-Solis et al., 2016; Whittle et al., 2015). Consistent with the metabolic changes, we found that GABRA5^LHA^ inhibition led to a significant decrease in interscapular BAT (iBAT) mRNA levels of the uncoupling protein *Ucp1*, *Cidea* and *Dio2*, but not *Prdm16*, *Pgc-1*α (Figure 2O). There was a significant change in the level of β-adrenergic receptor 3 (*Adrb3*) in inguinal WAT (iWAT) (Figure 2P). These genes are important regulators of thermogenesis, browning and beiging of adipose tissue, and lipolysis (Cao et al., 2011; Orozco-Solis *et al*., 2016; Whittle *et al*., 2015). Overall, these results suggest that chemogenetic inhibition of GABRA5^LHA^ suppresses whole body energy expenditure with a reduced expression of genes regulating thermogenesis and lipolysis.

**Figure 2.**
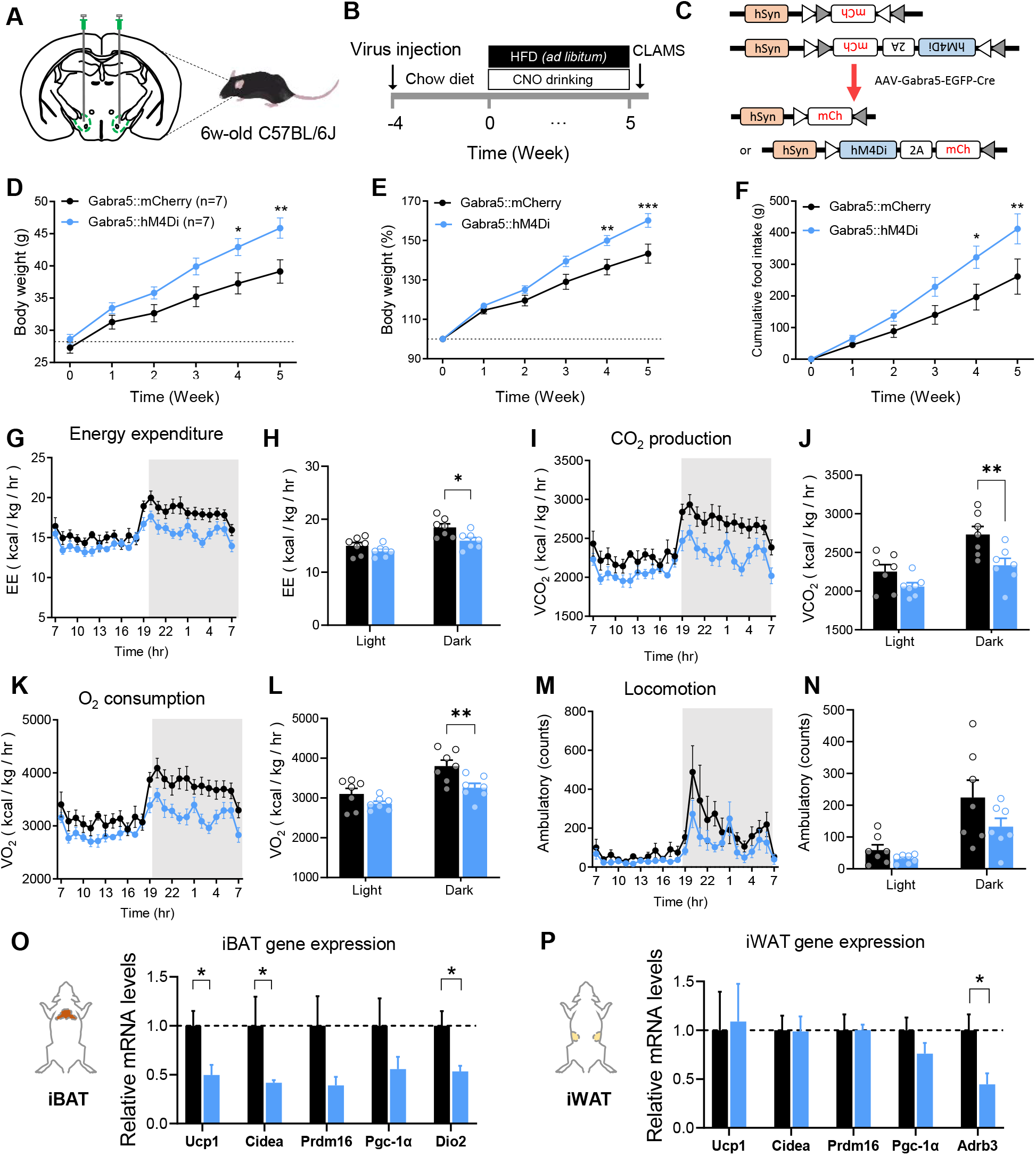
Chemogenetic inhibition of GABRA5^LHA^ suppresses energy expenditure. (A and B) Experimental schema and timeline for GABRA5^LHA^ infected with inhibitory DREADD hM4Di. (C) Schematic diagram of Gabra5::mCherry (top) and Gabra5::hM4Di (bottom) viruses with AAV-Gabra5-EGFP-Cre virus. (D and E) Body weight on HFD with CNO drinking in gram (D) and in percentage (E). n = 7 mice per group. (F) Cumulative food consumption on HFD with CNO drinking. (G and H) Real-time monitoring curve of energy expenditure and quantification of energy expenditure in light and dark cycle. (I and J) Real-time monitoring curve and quantification of carbon dioxide production in light and dark cycle. (K and L) Real-time monitoring curve and quantification of oxygen consumption in light and dark cycle. (M and N) Real-time monitoring curve and quantification of locomotor activity in light and dark cycle. Two-way ANOVA comparing Gabra5::mCherry and Gabra5::hM4Di with CNO drinking (n = 7 mice per group). (O) Molecular profiling of iBAT after chemogenetic inhibition of GABRA5^HA^. Genes tested: *Ucp1* (p < 0.05), *Cidea* (p > 0.05), *Prdm16* (p > 0.05), *Pgc-1*α (p > 0.05), *Dio2* (p < 0.05). Unpaired t tests comparing treatments (n = 6 samples per group). (P) Molecular profiling of iWAT after chemogenetic inhibition of GABRA5^LHA^. Genes tested: *Ucp1* (p > 0.05), *Cidea* (p > 0.05), *Prdm16* (p > 0.05), *Pgc-1*α (p > 0.05), *Adrb3* (p < 0.05). n = 6 samples per group. Data represents Mean ± SEM. *, p<0.05; **, p<0.01; ***, p<0.001.

### Gene-silencing of Gabra5 in LHA reduces obesity

To assess whether Gabra5 regulates body weight and food intake, we employed gene-silencing of Gabra5 in LHA of HFD mice. We developed (Figures S2A-S2C) and injected lentivirus carrying mouse *Gabra5*-specific short hairpin RNA (shRNA) into LHA (Figures 3A-3C), whose knockdown efficiency was confirmed using immunohistochemistry (Figures S2D-S2F). Gene-silencing of Gabra5 (shGabra5 group) showed a significant decrease in body weight after 5 weeks of injection (Figure 3D), without affecting food intake compare to the control (Scrambled group) (Figure 3F). To evaluate the changes of each peripheral organ with gene-silencing, we dissected key organs (Figure 3E) and observed the weights of iWAT and perigonadal WAT (pWAT) significantly decreased in shGabra5 mice (Figure 3G). In contrast, the weights of liver, kidney, spleen, heart and quadriceps were not different in shGabra5 mice compared to Scrambled mice (Figures 3E and 3G). BAT was slightly reduced in shGabra5 mice, although it was not significantly different (Figure 3H). To investigate the changes at the cellular level, we stained the adipose tissues with hematoxylin and eosin (H&E). The adipocyte size of iWAT and pWAT were significantly reduced in shGabra5 mice (Figures 3I-3L). BAT histology also revealed significantly smaller lipid droplets of adipocytes in shGabra5 mice (Figures 3M and 3N). To define hepatic morphological alteration caused by gene-silencing at the cellular level, we also observed hepatic cells using H&E staining. There was no significant change of lipid droplet in the hepatic cells (Figures 3O and 3P). Taken together, these results indicate that gene-silencing of Gabra5 prevents adipose tissue-specific weight increases with HFD.

**Figure 3.**
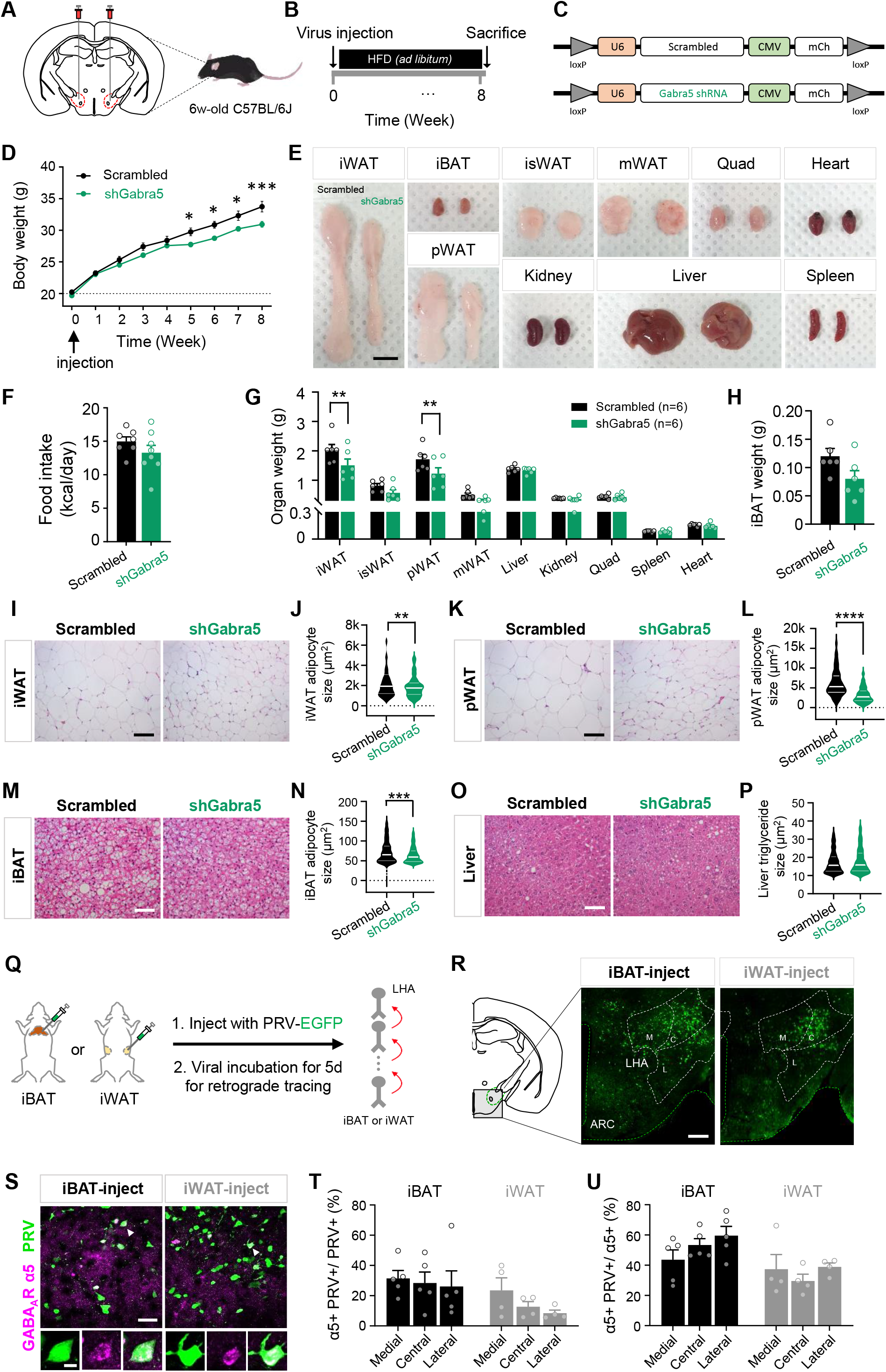
Knockdown of Gabra5 in LHA, innervating iBAT and iWAT, reduces obesity. (A and B) Experimental schema and timeline for LHA injection. (C) Schematic diagram of scrambled (top) and shGabra5 (bottom) viruses. (D) Curves of bodyweight in scrambled and shGabra5 mice. Knockdown of Gabra5 in LHA reduces body weight in grams (n = 7-8 per group). (F) Average food intake per day in scrambled and shGabra5 mice. (E and G) Quantification of organ weight and representative images of each organ between scrambled and shGabra5 group (n = 6 per group). Scale bar, 1 cm. (H) Average weight of iBAT. (I and J) H&E of inguinal WAT (I) and quantified adipose size of iWAT (J) of scrambled and shGabra5 group (n = 6 mice per group). Scale bar, 100 μm. n=180, 262 cells, respectively. (K and L) H&E of perigonadal WAT (L), and quantified adipose size of pWAT (L) of scrambled and shGabra5 group (n = 6 mice per group). Scale bar, 100 μm. Scale bar, 100 μm. n=120, 180 cells, respectively. (M and N) H&E of interscapular BAT (M) and quantified adipocyte size of iBAT (N) of scrambled and shGabra5 group (n = 6 mice per group). Scale bar, 100 μm. Scale bar, 100 μm. n=392, 391 cells, respectively. (O and P) H&E of liver (O) and quantified triglyceride size of liver (P) of scrambled and shGabra5 group (n = 6 mice per group). Scale bar, 100 μm. n=732, 725 cells, respectively. (Q) Schema for identifying LHA neurons projecting polysynaptically to iBAT and iWAT. (R) Confocal images for PRV-infected cells in LHA at 5 days post-infection of PRV in iBAT and iWAT. Bregma, −1.58 mm AP. Scale bar, 100 μm. (S) Gabra5 is colored by magenta, and PRV-EGFP is colored by green. Top, representative IHC with inset of region. Scale bar, 50 μm. Bottom, magnification of ROI in LHA regions. Scale bar, 10 μm. (T and U) Quantification between Gabra5 and PRV retrogradely labeled from iBAT and iWAT 5 days post-injection (n =3-4 mice per group). Data represents Mean ± SEM. *, p<0.05; **, p<0.01; ***, p<0.001.

### GABRA5^LHA^ polysynaptically project to BAT and WAT

To determine whether GABRA5^LHA^ project to adipose tissues, we infected iBAT and iWAT with a recombinant pseudo rabies virus (PRV) that enables retrograde tracing of polysynaptically connected circuit (Bartness et al., 2005; Ryu et al., 2017; Schneeberger et al., 2019). The mice were sacrificed 5 days after injection in iBAT and iWAT (Figure 3Q). In previous studies, several brain regions were found to project to iBAT and iWAT, including primary sensory cortex, paraventricular hypothalamus (PVH), periaqueductal gray (PAG), LHA and raphe pallidus nucleus (RPa) (Ryu et al., 2015; Ryu *et al*., 2017; Schneeberger *et al*., 2019; You et al., 2020). Among these areas, we found that iBAT- or iWAT-projecting neurons were detected in the medial, central and lateral part of the caudal LHA (Figures 3R and S3A). Then, we performed immunohistochemistry to test whether iBAT- or iWAT-projecting neurons (EGFP-positive) overlap with Gabra5-positive neurons within LHA. Interestingly, we found that ∼30% of the iBAT-projecting neurons and ∼15% of the iWAT-projecting neurons overlapped with Gabra5 throughout LHA (Figures 3S-3U). These results imply that GABRA5^LHA^ are responsible for polysynaptically innervating iBAT and iWAT, which was further supported by the results from the tissue clearing and light-sheet microscopic images (Figures S3B and S3C, Supplemental Video 1).

### Reactive astrocytes in LHA in response to HFD

Reactive astrocytes are observed in several regions of hypothalamus of HFD mice, such as arcuate nucleus (ARC), medial preoptic, paraventricular and dorsomedial hypothalamus (Buckman *et al*., 2013). Unlike other hypothalamic regions, reactive astrocytes in LHA are not defined yet. To define the reactivity of astrocytes in LHA, we performed immunohistochemistry to examine the expression of astrocyte markers in LHA after 20 weeks of HFD feeding (Figures 4A). As a result, astrocytes showed significantly hypertrophied signals in GFAP and S100β (Figures 4B-4F). Volume of GFAP-positive astrocytes significantly increased after 3D-rendering in HFD mice (Figures 4D and 4G). Sholl analysis of individual reactive astrocytes (Figure 4H) showed that the summation of intersects (Figures 4I and 4J), ramification index (Figure 4K) and ending radius (Figure 4L), which is an indicator of astrocytic territory, increased significantly in HFD compared to control. Therefore, these results indicate that the astrocytes in LHA become reactive in response to HFD as evidenced by the prominent morphological hypertrophy.

**Figure 4.**
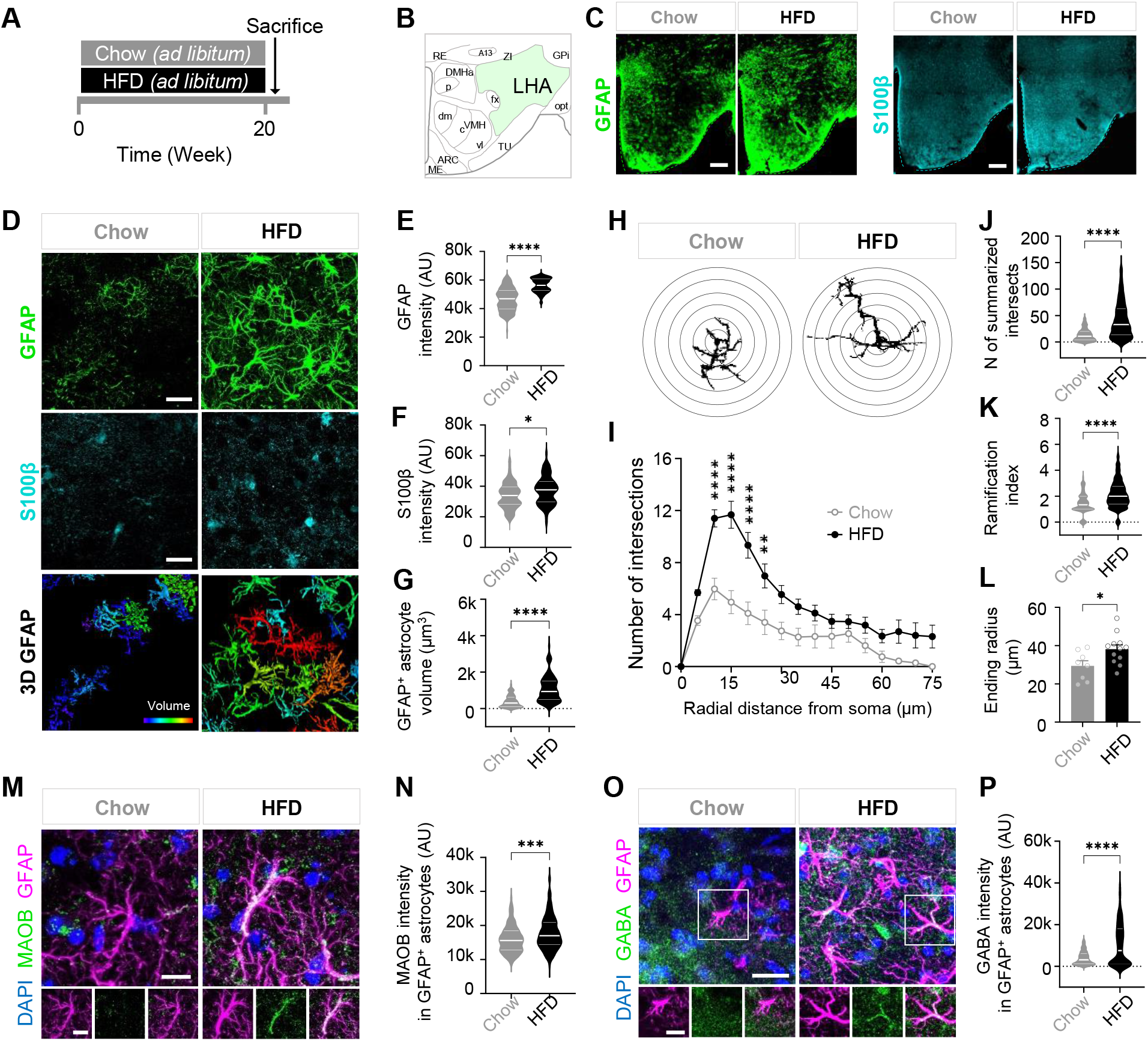
Astrocytes in LHA show hypertrophy in response to HFD. (A) Timeline of two different diet; chow diet and high fat diet group. (B and C) Coronal images of GFAP and s100β images in chow and HFD mouse in LHA. GFAP is colored by green and s100β is colored by cyan. Scale bar, 100 μm. (D) Significantly increased GFAP, s100β and 3D GFAP after HFD. Scale bar, 20 μm. (E - G) Violin plots showing quantification of GFAP intensity (E), S100β intensity (F) and GFAP-positive astrocyte volume (G) in chow and HFD mice. Unpaired t test comparing treatments (n = 4 - 6 mice per group). (H) Representative image for Sholl analysis of an astrocyte in the LHA from the GFAP-stained image in D. The interval of the concentric circles is 1μm. (I) The number of process intersections was significantly increased in GFAP^+^ astrocytes in the LHA of chow and HFD mice. (J and K) Number of summarized intersections (J) and ramification index (K) of GFAP^+^ astrocytes in chow and HFD mice. n= 381 - 411 cells per group, n=4 - 6 mice per group. (L) The ending radius of GFAP+ astrocytes in LHA of chow and HFD mice. All data are the average of cells per slice; 2 slices per mouse. n= 4 - 6 mice per group. (M) Immunostaining for MAOB and GFAP in LHA of chow and HFD mice. Top, Scale bar, 10 μm. Bottom, magnification of ROI in LHA regions. Scale bar, 10 μm. (N) Quantification of MAOB intensity in GFAP^+^ astrocytes. n = 152 cells per group. n= 4 - 6 mice per group. (O) Immunostaining for GABA and GFAP in LHA of chow and HFD mice. Top, Scale bar, 20 μm. Bottom, magnification of ROI in LHA regions. Scale bar, 10 μm. (P) Quantification of GABA intensity in GFAP^+^ astrocytes. n = 338-367 cells per group. n= 4 - 6 mice per group. Data represents Mean ± SEM. *, p<0.05; **, p < 0.01; ***, p < 0.001; ****, p < 0.0001.

We have previously reported that Aβ plaques cause an increase in the activity of astrocytic MAOB, which has been shown to produce GABA leading to a decreased neuronal activity in animal models of AD model (Jo *et al*., 2014). On the basis of this previous report, we hypothesized that those reactive astrocytes in HFD mice would have high levels of MAOB and GABA. We performed immunostaining in HFD mice and found that astrocytic MAOB signals were significantly increased (Figures 4M and 4N). We also observed that astrocytic GABA in HFD mice was significantly elevated (Figures 4O and 4P), with no change in neuronal Gabra5 level (Figures S3D and S3E).

Taken together, these results indicate that MAOB-mediated GABA in reactive astrocytes is significantly increased in LHA of HFD mice.

### Gene-silencing of astrocytic MAOB in LHA prevents obesity

We then hypothesized whether astrocyte-specific gene-silencing of MAOB in LHA can prevent DIO. To establish astrocyte-specific knockdown of MAOB, we used cre-dependent pSico-shMAOB with astrocyte-specific GFAP-cre viruses in LHA (shMAOB group) and cre-dependent pSico-Scrambled virus as control (Scrambled group) (Figures 5A and 5C). Their knockdown efficiency was confirmed by immunohistochemistry in LHA (Figures S4A-S4C). Both groups were fed with HFD (Figure 5B) and we found a significant reduction of astrocytic GABA in shMAOB group (Figures 5D and 5E). In addition, shMAOB mice showed a significant decrease in body weight after 8 weeks of HFD feeding without affecting food intake (Figures 5F and 5G). We next analyzed and compared the weights of each organ between shMAOB and Scrambled mice and found that shMAOB mice exhibited a significant reduction of weight in iWAT, pWAT and BAT (Figures 5H-5J). iWAT (Figures 5K and 5L), pWAT (Figures 5M and 5N) and BAT (Figures 5O and 5P) histology revealed reduced lipid droplets in shMAOB mice compared to the Scrambled mice with no significant change in the liver (Figures 5Q and 5R). Taken together, our results indicate that gene-silencing of LHA-specific astrocytic MAOB reduces adipose tissue-specific weight thereby reducing body weight without compromising appetite.

**Figure 5.**
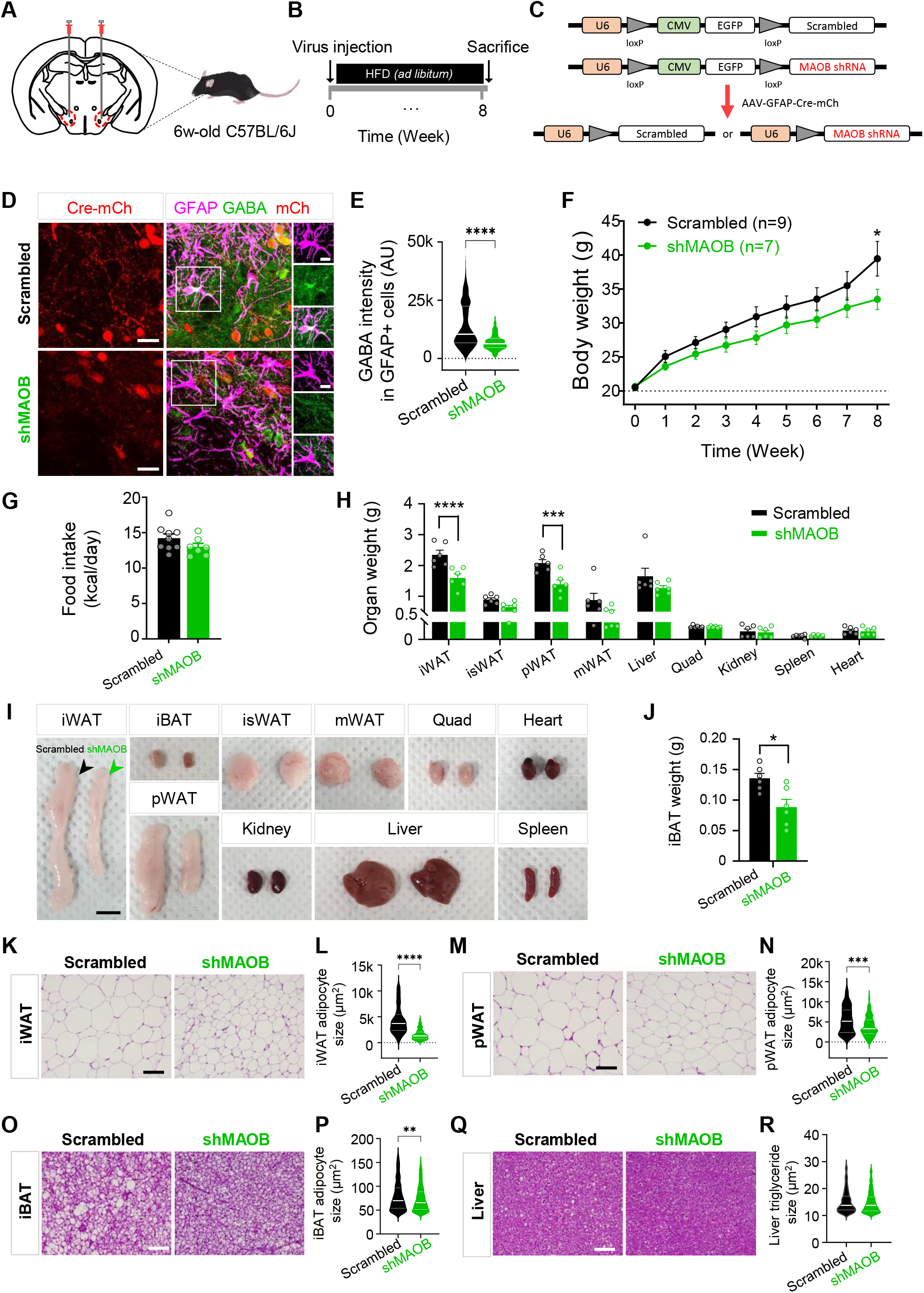
Gene-silencing of astrocytic MAOB in LHA prevents obesity. (A and B) Experimental schema and timeline for LHA injection. (C) Schematic diagram of Scrambled (top) and shMAOB (bottom) viruses with GFAP-cre virus. (D and E) Representative immunostaining for Cre (mch), GFAP and GABA in LHA (D) and quantification of GABA intensity in GFAP+ astrocytes between scrambled and shMAOB mice. Scale bar, 20 μm. Scale bar, 10 μm. (F) Inhibition of MAOB targeting LHA reduces body weight in grams (n = 7-8 per group). (G) No difference is observed in average food intake per day. (H and I) Quantification of organ weight and representative images of each organ between scrambled and shMAOB group (n = 6 per group). Scale bar, 1 cm. (J) Average weight of iBAT in scrambled and shMAOB group. (K and L) H&E of inguinal WAT (K) and quantified adipose size of iWAT (L) of scrambled and shMAOB group (n= 6 mice per group). Scale bar, 100 μm. n= 138, 155 cells, respectively. (M and N) H&E of perigonadal WAT (M) and quantified adipose size of pWAT (N) of scrambled and shMAOB group (n= 6 mice per group). Scale bar, 100 μm. n= 114, 147 cells, respectively. (O and P) H&E of iBAT (O) and quantified adipose size of iBAT (P) of scrambled and shMAOB group (n= 6 mice per group). Scale bar, 100 μm. n= 580, 641 cells, respectively. (Q and R) H&E of liver (Q) and quantified triglyceride size (R) of scrambled and shMAOB group (n= 6 mice per group). Scale bar, 100 μm. n=295, 569 cells, respectively. Data represents Mean ± SEM. *, p<0.05; **, p<0.01; ***, p<0.001; ****, p < 0.0001.

### Reducing GABA production via MAOB facilitates energy expenditure

Finally, we tested that inhibition of astrocytic GABA synthesis can decrease body weight. To test this hypothesis, we fed HFD to 6-week old C57BL/6J mice for 15 weeks till they reached near 50 g in bodyweight, after which they were treated with a recently developed a highly selective and reversible MAOB inhibitor, KDS2010 (Park et al., 2019) (Figures 6A and 6B). We measured body weight (Figures 6C and 6D) and food intake (Figure 6F) of chow, chow with KDS2010-treated, HFD and HFD with KDS2010-treated mice every week (Figure 6E). We observed a significantly potent decrease in body weight in KDS2010-treated HFD mice to the level of chow mice within 8 weeks (Figure 6E) without changing their food intake (Figure 6F). In contrast, an irreversible MAOB inhibitor Selegiline showed only a transient reduction in the body weight (Figures S5A-S5D). These effects appear to be mainly through MAOB inhibition in the brain rather than peripheral system (Figures S5E-S5I). We then examined the body composition of each group using EchoMRI for measurement of fat and lean mass. There was a significant reduction only in fat mass (Figure 6G), not in lean mass (Figure 6H). These results suggest that fat-specific weight loss is due to the MAOB inhibition by a reversible MAOB inhibitor in the brain.

**Figure 6.**
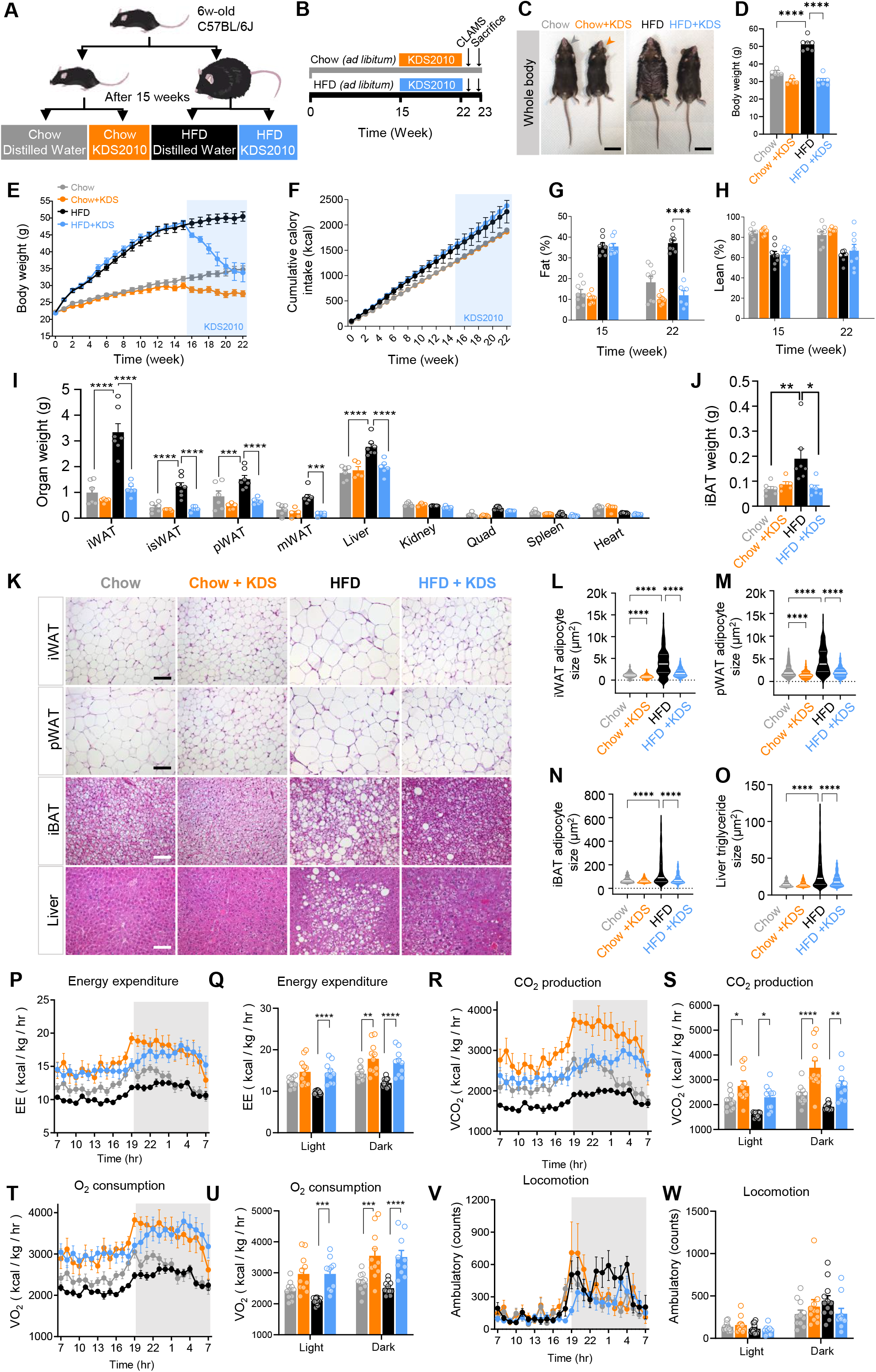
Reducing GABA production via MAOB reduces obesity. (A) Four groups of different diet with drug treatment; chow diet with primary distilled water, chow diet with KDS2010, high fat diet with primary distilled water and high fat diet with KDS2010 treatment group. (B) Experimental timeline. (C and D) Representative images of each group mouse before sacrifice. Average body weight of four groups (n= 5 – 7 mice per group). Scale bar, 3 cm. (E) Curves representing the kinetics of change in body weight among chow, chow with KDS2010, HFD, HFD with KDS2010 mice over the 22 weeks following HFD treatment. Light blue box means KDS2010 treatment. n= 8 mice per group. (F) Cumulative food intake in chow, chow with KDS2010, HFD, HFD with KDS2010 mice over the 22 weeks. n= 8 mice per group. (G and H) Quantification of percentage change of fat mass (G) and lean mass (H) at before (15 weeks) and after (22 weeks) KDS2010 treatment. (I and J) Quantification of organ weight (I) and average weight of iBAT (J) of chow, chow with KDS2010, HFD, HFD with KDS2010 group. (K – O) H&E of iWAT, pWAT, iBAT and liver (K). Scale bar, 100 μm. Quantified adipose size of iWAT (L), pWAT (M), iBAT (N) and liver (O) of chow, chow with KDS2010, HFD, HFD with KDS2010 group. (P - W) HFD with KDS2010 activate energy expenditure (P,Q), carbon dioxide production (R,S), oxygen consumption (T,U), but not in locomotor activity (V,W). n = 11, 11, 11 and 8 for respective group. Data represents Mean ± SEM. *, p<0.05, **, p < 0.01; ***, p < 0.001; ****, p < 0.0001.

Consistent with the fat mass reduction, we observed that the weights of iWAT, isWAT, pWAT and BAT in HFD with KDS2010-treated mice (Figure S6A) were significantly reduced to that of chow mice (Figures 6I and 6J). In addition to the reduction of fat mass, KDS2010 also reduced the weight of liver induced by long-term treatment of HFD (Figure 6I). Then, we examined the tissues at the cellular level using H&E staining. Consistent with the weight of each organ, histology of iWAT, pWAT, and BAT revealed a significantly reduced size of adipocytes in HFD with KDS2010-treated mice compared to HFD mice (Figures 6K-6N). Previous studies have shown that HFD induces nonalcoholic fatty liver disease (Recena Aydos et al., 2019). There was a significant increase in triglycerides in the liver in HFD mice, similar to the phenotype of fatty liver, which was significantly reduced in HFD with KDS2010-treated mice (Figure 6O). Taken together, pharmacological inhibition of MAOB with KDS2010 effectively and rapidly reduces obesity without affecting food intake.

To determine whether MAOB inhibition can promote energy expenditure, we measured metabolic parameters using CLAMS. HFD with KDS2010-treated mice have significantly higher energy expenditure (Figures 6P and 6Q), carbon dioxide production (Figures 6R and 6S) and oxygen consumption (Figures 6T and 6U) than HFD mice in both light and dark cycle with no significant difference in locomotor activity (Figures 6V and 6W). HFD with KDS2010-treated mice also showed significantly improved glucose tolerance (Figures S6B and S6C) and insulin sensitivity compared to HFD mice (Figures S6D and S6E). Overall, these results imply that pharmacological inhibition of MAOB facilitates energy expenditure, leading to bodyweight decrease.

### A reversible MAOB inhibitor reduces tonic GABA in LHA

We observed that GABRA5^LHA^ pacemaker firing is suppressed by GABA-meditated inhibition in HFD mice (Figures 1N and 1O). The mode of GABA action can be either phasic or tonic inhibition (Bhattarai et al., 2011; Farrant and Nusser, 2005). So, we asked whether GABRA5^LHA^ is either phasically or tonically inhibited in HFD mice. To test this possibility, we performed whole-cell patch-clamp recordings in the acutely prepared LHA slices (Figure 7A). We assessed GABA_A_ receptor-mediated phasic and tonic GABA currents by measuring the baseline current shift upon GABA_A_R antagonist, bicuculline, in the presence of ionotropic glutamate receptor antagonists, APV (50μM) and CNQX (20μM), as described previously (Jo *et al*., 2014; Lee et al., 2010). HFD mice showed a significant increase of tonic GABA current compared to chow mice which was reduced by KDS2010 treatment (Figure 7B-7F). There was no significant difference in GABA-induced full activation current which was induced by 10 μM GABA (Figure 7G). The amplitude and frequency of spontaneous inhibitory post-synaptic currents (sIPSCs) were not significantly altered, indicating that phasic or synaptic GABA was not altered (Figure 7H and 7I). The capacitance of LHA neurons was not affected by HFD (Figure 7J). These results indicate that GABA action for GABRA5^LHA^ is mediated by tonic inhibition and reversible KDS2010 significantly attenuated the tonic inhibition.

**Figure 7.**
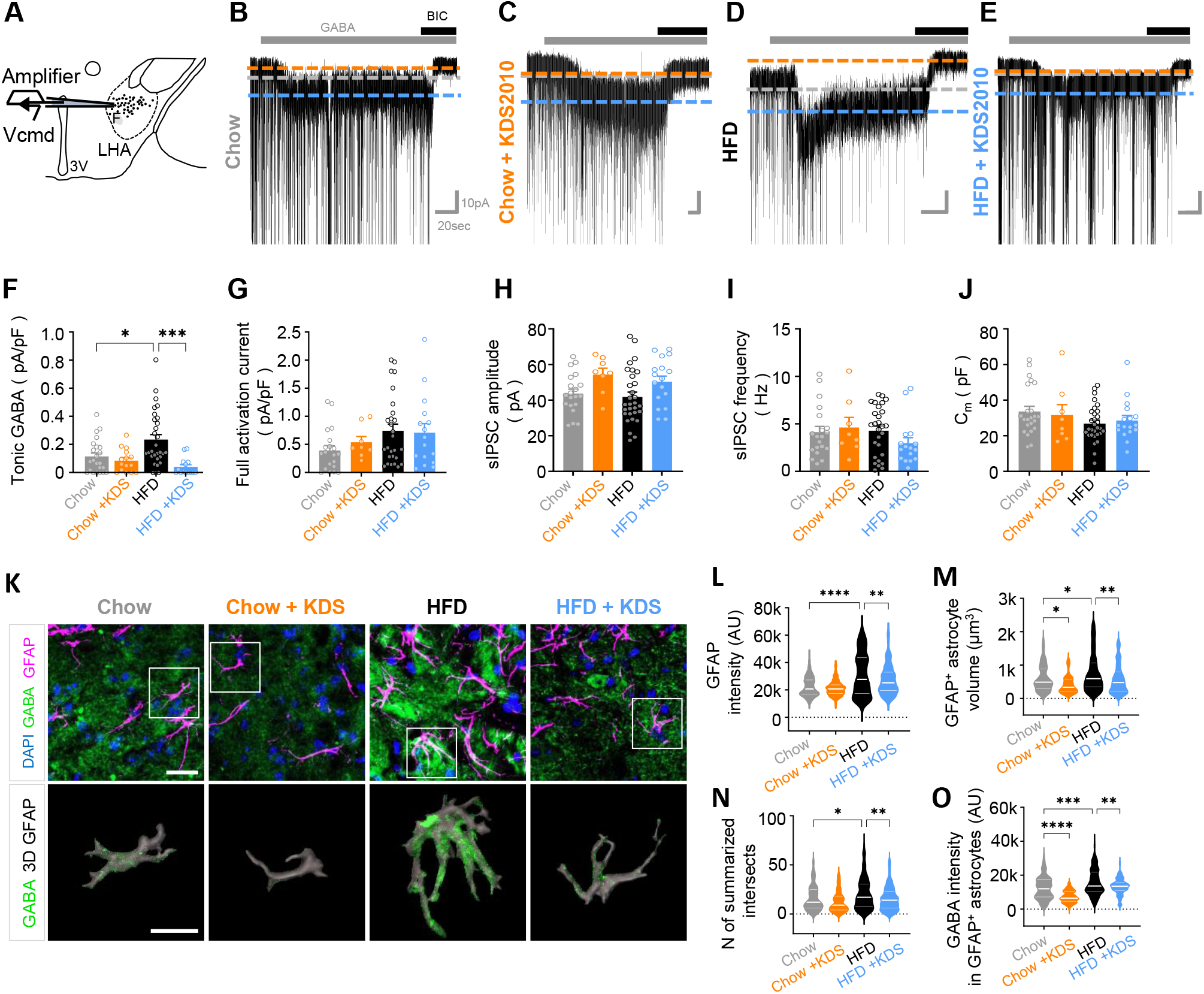
A reversible MAOB inhibitor reduces tonic GABA in LHA. (A) Schema of whole-cell patch-clamp recording in LHA neurons. (B - E) Representative traces of GABA_A_ receptor-mediated currents recorded from chow diet (B), chow diet with KDS2010 (C), high fat diet (D), high fat diet with KDS2010 (E) group. Each dot represents one cell (n = 6, 3, 8, and 3 mice, respectively). 50uM of BIC, 10uM of GABA were applied. (F) GABA_A_ receptor-mediated tonic GABA current measured from LHA neurons in each group. (G) GABA-induced full activation current measured from LHA neurons in each group. (H and I) Frequency and amplitude of sIPSC measured from LHA neurons in each group. (J) Capacitance of LHA neurons in each group. (K) Top, Immunostaining for GABA and GFAP in LHA of chow, chow with KDS2010, HFD and HFD with KDS2010 group. Scale bar, 20 μm. Bottom, magnification of ROI. 3D rendering GFAP with GABA immunostaining. Scale bar, 10 μm. (L) Quantification of GFAP intensity in GFAP^+^ astrocytes. n = 438, 222, 294 and 112 cells per group. n= 4 - 6 mice per group. (M) Quantification of volume of GFAP^+^ astrocytes. n = 89, 69, 179 and 117 cells per group. n= 4 - 6 mice per group. (N) Number of summarized intersections of GFAP^+^ astrocytes in each group. n= 128, 94, 169 and 205 cells per group, n=4 - 6 mice per group. (O) Quantification of GABA intensity in GFAP^+^ astrocytes. n = 104, 91, 160 and 113 cells per group. n= 4 - 6 mice, respectively. Data represents Mean ± SEM. *, p<0.05, **, p < 0.01; ***, p < 0.001; ****, p < 0.0001.

To examine the reactivity of astrocytes and astrocytic GABA level after KDS2010 treatment, we performed immunostaining in LHA. We found that GFAP signals and GFAP-positive astrocyte volume were significantly reduced in LHA of HFD with KDS2010-treated mice compared to HFD mice (Figures 7K-7M). We also observed that the summation of intersects was significantly reduced in HFD with KDS2010-treated mice (Figure 7N). As expected, we found that MAOB expression in astrocyte was significantly reduced in LHA of HFD with KDS2010-treated mice compared to HFD mice (Figures S6F and S6G). To test whether inhibition of MAOB reduces the GABA level in astrocytes, we performed immunostaining and found that astrocytic GABA signals were significantly reduced in HFD with KDS2010-treated mice compared to that of HFD mice (Figure 7O). Taken together, these results indicate that MAOB inhibition by the reversible inhibitor KDS2010 reduces astrocytic reactivity and GABA level in LHA, thereby attenuating tonic inhibition of LHA neurons in HFD mice.

## DISCUSSION

In the present study, we have discovered the existence of a unique population of energy expenditure-facilitating neurons in LHA. These neurons are uniquely expressing the high-affinity extrasynaptic GABA_A_ containing Gabra5 subunit and at the same time GABA-producing, *i.e*., GABAergic GABRA5^LHA^. Although GABAergic, these neurons appear to be projection neurons, projecting polysynaptically to iBAT and iWAT. These neurons show unique electrical properties of low-threshold spiking and pacemaker firing at a firing frequency of around 5 Hz. The presence of Gabra5 allows sensing of tonically released low concentration of extrasynaptic GABA, which has been recently characterized to be synthesized by MAOB and released by reactive astrocytes under pathological conditions such as in Alzheimer’s disease and Parkinson’s disease (Jo *et al*., 2014; Park *et al*., 2019). We have also demonstrated for the first time the causal relationship between reactive astrocytes and thermogenesis/fat storage in DIO mouse model via the complex GABA signaling in LHA, composed of increase in astrocytic tonic GABA, activation of neuronal GABA_A_ receptors containing α5 subunit, reduction in pacemaker firing in GABRA5^LHA^, attenuation of thermogenesis and augmentation of fat storage in peripheral adipose tissues (Figure S7A). These cascade of events in DIO model can be mimicked by activation of Gi-DREADD expressed in GABRA5^LHA^ and reversed by pharmacological inhibition or gene-silencing of MAOB to reduce astrocytic GABA and gene-silencing of Gabra5 to reduce tonic GABA inhibition in GABRA5^LHA^ (Figure S7B). Strikingly, this reversal of DIO can be achieved without compromising appetite. Our study proposes the pathological role of glia-neuron interaction via astrocytic GABA and neuronal Gabra5 and its contribution to thermogenesis/fat storage and DIO with minimal contribution to food intake.

### GABAergic GABRA5^LHA^ innervate BAT or WAT

Recent research highlights that LHA lacks inhibitory interneurons with locally ramifying axons (Burdakov and Karnani, 2020). This suggests that the GABAergic neurons in LHA project to outside of LHA rather than locally inhibit the nearby neurons. LHA contains a large number of GABAergic neurons expressing GAD65, GAD67 and VGAT (Elias et al., 2008; Harthoorn et al., 2005; Jennings *et al*., 2013; Jennings et al., 2015; Karnani *et al*., 2013). These neurons are likely to be highly diverse and subdivided into many subpopulations with distinct characteristics and functions. In the current study we have identified that GABAergic GABRA5^LHA^ project to outside of LHA and polysynaptically innervate iBAT and iWAT. LHA regulates both thermogenesis and lipolysis in brown adipocytes through the sympathetic nervous system (SNS) (Bamshad et al., 1999; Bartness et al., 2010; Cannon and Nedergaard, 2004; Contreras *et al*., 2017). LHA also regulates lipolysis of iWAT through SNS (Bartness *et al*., 2010; Cannon and Nedergaard, 2004; Richard and Picard, 2011). It has been reported that in mice injected with PRV into iBAT, PRV-infected neurons in LHA overlap with MCH and orexin (Izawa et al., 2021). However, there is still a remaining undefined population that innervates iBAT and iWAT in LHA. We propose that the remaining population is in part GABAergic GABRA5^LHA^. In previous studies, Orexin neurons in LHA were shown to project to rostral raphe pallidus (rRPa) in the brainstem (Contreras *et al*., 2017; Tupone *et al*., 2011), which are known as sympathetic premotor neurons to control sympathetic output and activate iBAT (Morrison et al., 2014). Consistently, viral tracing evidence further demonstrates that sympathetic nerves that innervate subcutaneous WAT originate from rRPa (Nguyen et al., 2014), which then project to sympathetic preganglionic neurons in the spinal intermediolateral nucleus (IML) (Morrison et al., 2012). Our results from the tissue clearing and light-sheet microscopic images suggest that GABAergic Gabra5^LHA^ might project to PAG before reaching to rRPa (Figure S3C). We propose that GABAergic Gabra5^LHA^, just like orexin neurons in LHA, may also project to rRPa possibly after passing through PAG and innervate iBAT and iWAT through IML and sympathetic ganglion. Future investigations are needed to determine the exact polysynaptic circuits for GABAergic Gabra5^LHA^, projecting to iBAT and iWAT.

### Astrocytic GABA suppresses the activity of Gabra5^+^ neurons

Our study sheds a light on the missing link between the reactive astrogliosis and obesity, delineating the molecular and cellular mechanisms of how MAOB-dependent production of GABA leads to inhibition of energy expenditure and facilitation of fat storage. Elevated activity of MAOB and elevated levels of MAOB-dependent GABA have been highly implicated in the reactive astrocytes that are found in various neuroinflammatory diseases such as Alzheimer’s disease, Parkinson’s disease, stab-wound injury model and et cetera (Chun et al., 2021; Jo *et al*., 2014; Nam et al., 2021; Pandit et al., 2020; Shim et al., 2019). The growing list of reports all point to the common molecular mechanism of how resting astrocytes transform into reactive astrocytes via the putrescine-degradation pathway involving MAOB under the conditions of aversive stimulations such as toxin challenges and viral infections, which usually accompany neuroinflammation (Chun *et al*., 2018). In the current study, we have also discovered the same common molecular mechanism at work in LHA to cause the reactive astrogliosis and the production of GABA in MAOB-dependent fashion. Then, what is the toxin that turns on this mechanism in LHA? It has been reported that chronic HFD induces hypothalamic inflammation, which is associated with reactive astrocytes in the hypothalamus (Thaler *et al*., 2012), raising a possibility that the high fat nutrition itself could be the trigger for reactive astrogliosis and GABA production. Indeed, we have recently found that astrocytes in culture start to produce GABA when they are challenged with elevated levels of fatty acids (Lee et al., 2018). Consistent with our findings, it has been reported that after chronic overnutrition, extracellular neurotransmitters such as GABA levels become elevated in mediobasal of hypothalamus (Zhang *et al*., 2017). In this study, we have further demonstrated that the MAOB-dependent astrocytic tonic GABA induces a strong neuronal inhibition in GABRA5^LHA^. In addition, the MAOB-dependent H_2_O_2_ has been recently implicated in neurodegeneration and brain atrophy in Alzheimer’s disease (Chun et al., 2020b). Although we have not investigated further, we expect that the reactive astrocytes in LHA would produce excess amount of toxic H_2_O_2_ in MAOB dependent fashion, further exacerbating the reactive astrogliosis. The excess amount of toxic H_2_O_2_ would cause neuronal death of the neighboring neurons in LHA under chronic obesity condition. Indeed, consumption of dietary fats induce apoptosis of neurons and a reduction of synaptic inputs in LHA (Moraes *et al*., 2009). This possibility of H_2_O_2_-induced neurodegeneration in LHA awaits future investigation.

### KDS2010, a reversible MAOB inhibitor, as a new therapeutic strategy for anti-obesity drug development

Our study add a new dimension to the existing anti-obesity drugs. It has been reported in previous studies that drugs for weight loss have a significant history of safety risks, including cardiovascular and psychiatric complications (Cheung et al., 2013; Kim et al., 2014a). Most of the obesity drugs that target neurons in the hypothalamus are known to suppress appetite rather than to increase energy expenditure (Cheung *et al*., 2013). Based on our study, we propose that selective inhibition of MAOB may be potential molecular targets for treating obesity to overcome the limitations of neuron-target obesity drugs. By using three pharmacological inhibitors, KDS2010, KDS1524, and selegiline, with differential properties, we have gained useful insights about designing effective therapeutic strategies. By comparing the irreversible inhibitor selegiline and reversible inhibitor KDS2010, we discovered that KDS2010 showed long-lasting effects compared to selegiline, implying that reversibility of MAOB inhibitor is critical for long-lasting efficacy. These results are consistent with our previous reports on the superior effect of reversible MAOB inhibitors on animal models of Alzheimer’s disease and Parkinson’s disease (Nam *et al*., 2021; Park *et al*., 2019). Furthermore, by comparing the BBB-permeable KDS2010 and the less BBB-permeable KDS1524, we discovered that KDS2010 showed a far superior effect than KDS1524, implying that the central MAOB in the brain is a far more effective target for developing anti-obesity drug than the peripheral MAOB. The effect of inhibiting MAOB in the brain can also affect other hypothalamic regions, such as ARC and PVH, where reactive astrocytes are readily observed after HFD (Buckman *et al*., 2013). This could explain why KDS2010 treatment showed a much steeper weight loss than the LHA-specific MAOB gene-silencing. KDS2010 treatment should ameliorate the aberrant reactive astrogliosis and tonic GABA inhibition throughout the various hypothalamic regions, thus eliminating the undesirable inhibition of neuronal activity. It would be interesting to investigate the existence and physiological roles of Gabra5-positive neurons in other hypothalamic regions. The difference in the long-term efficacy between selegiline and KDS2010 has been mechanistically explained by how reversible and irreversible inhibitors differentially act on the MAOB enzyme; irreversible inhibitors like selegiline covalently modify the MAOB enzyme and destroy the enzyme itself to turn-on the compensatory mechanism of DAO, whereas reversible inhibitors occupy the active site of MAOB competitively, resulting in an intact MAOB enzyme with no compensatory mechanism (Park *et al*., 2019). More importantly, selective inhibition of MAOB does not affect food intake and appetite. These novel features of MAOB inhibitors will help develop better anti-obesity drugs in the future.

In summary, we propose that GABRA5^LHA^ are distinct GABAergic-projecting and pacemaker-firing neurons which facilitate energy expenditure through adipocyte tissues. These findings establish the GABRA5^LHA^ as key players modulating GABA via astrocyte-neuron interaction in hypothalamus of DIO mouse model. Our study raises new molecular targets to combat obesity without compromising appetite.

## Author contributions

M Sa, ES Yoo, W Koh, MG Park, HJ Jang, YR Yang, J Lim, W Won, J Kwon, M Bhalla, H An, Y Seong performed experiments. KD Park, PG Suh, JW Sohn, CJ Lee supervised the analysis. M Sa and CJ Lee wrote the manuscript.

## Acknowledgements

This work was supported by the Institute for Basic Science (IBS), Center for Cognition and Sociality (IBS-R001-D2). This study was also supported by the National Research Foundation (NRF) Grants from the Korean Ministry of Education, Science and Technology (2018M3C7A1056894, NRF- 2020M3E5D9079742) and KIST Grants (2E30954 and 2E30962).

## Declaration of interests

The authors declare that there is no conflict of interests.

## STAR Methods

### RESOURCE AVAILABILITY

#### Lead contact

Further information and requests for resources and reagents should be directed to and will be fulfilled by the lead contact Dr. C Justin Lee

#### Materials availability

The sequences of the shRNAs used in this study have been provided in the Supplementary figures. The viruses used in this study were provided by and are available with the Institute for Basic Science Virus Facility (https://www.ibs.re.kr/virusfacility) and Korea Institute of Science and Technology Virus Facility upon request (http://virus.kist.re.kr).

#### Data and code availability

Accession number is listed in the Key Resources Table. Microscopy data reported in this paper will be shared by the lead contact upon request. This paper does not report any original code. Any additional information required to reanalyze the data reported in this paper is available from the lead contact upon request.

### EXPERIMENTAL MODEL AND SUBJECT DETAILS

#### Animals and housing

All animal experiments were performed according to procedures approved by the Institutional Animal Care and Use Committee of IBS (Daejeon, South Korea) and Korea Institute of Science and Technology (Seoul, South Korea). All mice were maintained in a specific pathogen-free animal facility under a 12-h light-dark cycle (lights on at 8:00 AM) at a temperature of 21°C and allowed free access to water and food. All experiments performed on diet-induced obesity (DIO) mouse model were performed on C57BL/6J background were used originated from Jackson Laboratory (USA, stock number 000664). 6-week-old male C57BL/6J mice (DBL, Chungbuk, Republic of Korea) were fed a HFD (60% kcal fat, D12492, Research Diets Inc.) or chow (Teklad, 2018S, Envigo) for 6∼23 weeks. All experiments were done with age-matched controls.

### METHOD DETAILS

#### Stereotaxic injection

Mice were anesthetized using isoflurane anesthesia (induction: 3%–4%, maintenance: 1.5%–2%) and placed into stereotaxic frames (Kopf). The scalp was incised and a hole was drilled into the skull above the LHA (anterior/posterior, −1.58 mm; medial/lateral, −1.0 or +1.0 mm from bregma). Coordinates were identified using the Allen mouse brain atlas. For characterization studies, C57BL/6J mice were injected with 1.0 μL of AAV-mGabra5-EGFP-cre virus on both sides of LHA. For Gabra5 knockdown studies, C57BL/6J mice were injected with 1.0 μL of Lenti-pSicoR-Gabra5 shRNA-mCherry or Lenti-Scrambled-mCherry virus on both sides of LHA. For chemogenetic studies, AAV5-mGabra5-EGFP-Cre with AAV5-hSyn-DIO-hM4Di-mCherry (inhibition) or AAV5-hSyn-DIO-mCherry (control virus) were injected on both sides of LHA. For astrocyte-specific MAOB knockdown studies, AAV-GFAP-Cre-mCherry with Lenti-pSico-Scrambled-GFP or Lenti-pSico-shMAOB-GFP virus were used on both sides of LHA. The virus was loaded into a stainless needle and injected bilaterally into the LHA (dorsal/ventral, −5.0 mm) at a rate of 0.1 μlmin^−1^ for 10 min using a syringe pump (KD Scientific). At the end of the infusion, the needle was left in the brain for another 10 min to reduce backflow of the virus. Shortly after surgery, mice were translocated to their home cages.

#### Slice preparation for electrophysiology

Mice were deeply anaesthetized with vaporized isoflurane and then decapitated to isolate the brain. The isolated brains were quickly excised from the skull and submerged in ice-cold NMDG recovery solution containing: 93 mM of NMDG, 93 mM of HCl, 30 mM of NaHCO_3_, 20 mM of HEPES, 25 mM Glucose, 5 mM sodium ascorbate, 2.5 mM KCl, 1.2 mM NaH_2_PO_4_ (pH 7.4.). All the solution was gassed with 95% O_2_ and 5% CO_2_. The brain was glued onto the stage of a vibrating microtome (Linear Slicer Pro7, D.S.K) and 250-μm-thick coronal slices were prepared. For stabilization, slices were incubated in room temperature for at least 1 h in extracellular aCSF solution containing 130 mM of NaCl, 3.5 mM of KCl, 24 mM of NaHCO_3_, 1.25 mM of NaH_2_PO_4_, 1.5 mM of CaCl_2_, 1.5 mM of MgCl_2_, and 10 mM of d-(+)-glucose, pH 7.4. and simultaneously equilibrated with 25℃. Slices were transferred to a recording chamber that was continuously perfused with aCSF solution.

#### Patch-clamp recording

For the characterization of GABRA5^LHA^ cells, electrophysiological experiments were conducted with reference to previous study (Karnani *et al*., 2013). Patch electrodes (4-8 MΩ) were filled with an intrapipette solution containing: 120 mM of potassium gluconate, 10 mM of KCl, 1 mM of MgCl_2_, 0.5 mM of EGTA and 40 mM of HEPES (pH 7.2 adjusted with KOH). Resting membrane potential (mV) was measured at I=0 soon after membrane rupture. Step current was injected in current clamp mode to measure maximum firing frequency (Hz), which is the reciprocal of the average of the first four peak intervals (ISI) calculated after the highest injected current before the occurrence of spike inactivation. Spike ratio was determined by calculating X/(Y/2) from the number of spikes before (500 ms window, Y) and after (250 ms window, X) hyperpolarization by negative current step. Classification of cells, which is late spiking for <0.5, regular spiking for between 0.5 and 1.5, low-threshold spiking for >1.5 followed previous study (Karnani *et al*., 2013). For the measurement of spontaneous spike activity in GABRA5^LHA^, cell-attached patch was conducted as previously described (Heo et al., 2020). Patch electrodes (4-8 MΩ) were filled with normal aCSF solution. The slice chamber was mounted on the stage of an upright microscope and viewed with a 60× water immersion objective (numerical aperture = 0.90) with infrared differential interference contrast optics. Cellular morphology was visualized by a complementary metal oxide semiconductor camera and the Imaging Workbench software (INDEC BioSystems, ver. 9.0.4.0.).

#### Tonic GABA recording

Whole-cell patch-clamp recording was conducted as previously described (Jo *et al*., 2014). The holding potential was −60 mV. Pipette resistance was typically 6–8 MΩ and the pipette was filled with an internal solution consisting of: 135 mM of CsCl, 4 mM of NaCl, 0.5 mM of CaCl_2_, 10 mM of HEPES, 5 mM of EGTA, 2 mM of Mg-ATP, 0.5 mM of Na_2_-GTP, and 10 mM of QX-314, pH-adjusted to 7.2 with CsOH (278–285 mOsmol). Before measuring the tonic current, the baseline current was stabilized with D-AP5 (50 μM) and CNQX (20 μM) to isolate GABA_A_ receptor current from AMPAR and NMDAR. Electrical signals were digitized and sampled at 10-ms intervals with Digidata 1550 data acquisition system and the Multiclamp 700B Amplifier (Molecular Devices) using the pClamp10.2 software. Data were filtered at 2 kHz. The amplitude of the tonic GABA current was measured by the baseline shift in response to the bath application of bicuculline (50 μM) using the Clampfit software (ver. 10.6.0.13.). The frequency and amplitude of spontaneous inhibitory postsynaptic currents before bicuculline administration was detected and measured by Mini Analysis (Synaptosoft, ver. 6.0.7.)

#### Immunohistochemistry

Mice were deeply anaesthetized with isoflurane and transcardially perfused with 0.9% saline followed by ice-cold 4% paraformaldehyde (PFA). Excised brains were postfixed overnight at 4°C and transferred to 30% sucrose for 48 hours and cut with a frozen microtome in coronal 30 μm sections. Brain sections were translocated into 24-well plates filled with blocking solution (0.3% Triton X-100, 3% Donkey Serum in 0.1M PBS). Primary antibodies were added to blocking solution at desired dilution and slices were incubated in a shaker at 4°C overnight. Primary antibodies for immunostaining were anti-Gabra5 (rabbit, 1:200), Orexin A (rabbit, 1:100), Orexin B (rabbit, 1:500), MCH (rabbit, 1:200), GABA (rabbit, 1:200), GFAP (chicken, 1:500), S100β (rabbit, 1:200), MAOB (mouse, 1:100), NeuN (mouse or guinea pig, 1:500) and c-FOS (rabbit, 1:500). Antibody details can be found in the Key Resources Table. Unbound antibodies were washed off using PBS, followed by corresponding secondary antibody incubation (in blocking solution) for 1 or 2 hours at room temperature. Unbound antibodies were washed with PBS and DAPI was added to PBS (1:1500 dilution) in the second step to visualize the nuclei of the cells. Sections were mounted with fluorescent mounting medium (Dako) and dried. Series of fluorescent images were obtained by Zeiss LSM900 confocal microscope using a 20x, 40x, or 63x objective. Z stack images were processed using the ZEN Digital Imaging for Light Microscopy blue system (Zeiss, ver. 3.2) and ImageJ (NIH, ver. 1.52s.) software.

#### Image quantification

Confocal microscopic images were obtained in order to quantify the number of colocalized cells and expression were analyzed using the ImageJ (NIH) program. Fluorescence intensities were calculated using the mean intensity value of each fluorescence pixels in the marker-positive area. The marker-positive area was defined by thresholding and is converted into a binary mask. The mean intensity of immunostained pixels in the binary mask was calculated. Sholl analysis was performed on serially stacked and maximally projected confocal images as previously described (Chun et al., 2020a). Confocal images of brain sections immunostained with GFAP antibody were used for Sholl analysis. The Sholl analysis plugin applied in IMARIS software (Version 9.0.1, Oxford Instruments) constructs serially concentric circles at 10μm intervals from the center of GFAP signal (soma) to the end of the most distal process of each astrocyte. The number of intercepts of GFAP-positive processes at each circle and the radius of the largest circle intercepting the astrocyte are analyzed. For measuring GFAP-positive volume of astrocyte, GFAP signals were reconstructed to a 3-dimensional (3D) object by 3D reconstruction using IMARIS software (Version 9.0.1, Oxford Instruments).

#### Tissue isolation and histological analysis

Mice were deeply anesthetized with isoflurane and organs were immediately isolated from the mice body. we isolated inguinal white adipose tissues (iWAT), interscapular white adipose tissue (isWAT), perigonadal white adipose tissue (pWAT), mesenteric white adipose tissue (mWAT), Liver, quadriceps muscle (Quad), kidney, spleen, and heart. After measuring the weight of each organ, iBAT, iWAT, pWAT and Liver tissues were fixed with 4% PFA (Sigma-Aldrich, St. 538 Louis, MO) for overnight and conducted further processes. Histological changes of lipid droplets were examined by hematoxylin and eosin (H&E) staining. As counterstain, Mayer’s hematoxylin was used for every slide. H&E images were obtained with Eclipse TI-E microscope and were analyzed using the ImageJ (NIH) program.

#### Tissue clearing and light-sheet imaging

Mice were transcardially perfused with ice-cold PBS and then with the SHIELD perfusion solution (Passive clearing, Lifecanvas, 500ml kit - Cat. No. PCK-500) as described previously (Park et al., 2018). Dissected brains were incubated in the same perfusion solution at 4 °C for 48 h. Tissues were then transferred to the SHIELD-OFF solution and incubated at 4 °C for 24 h. Following the SHIELD-OFF step, brains were placed in the SHIELD-ON solution and incubated at 37 °C for 24 h. SHIELD-fixed brains were cleared passively for a couple of weeks (10–14 d at 45 °C for a mouse brain hemisphere) in buffer solution. Delipidated tissues were incubated in Protos-based immersion media until the tissue became transparent without any visible haze at the tissue–medium interface. 3D light-sheet images were taken by Zeiss Light sheet Fluorescence microscopy (LSFM) 7.

#### Metabolic analysis

For metabolic analysis, mice were measured using the Comprehensive Lab Animal Monitoring System (CLAMS, Columbus Instruments). All mice for metabolic analysis were maintained in an animal facility under a 12-h light-dark cycle (lights on at 7:00 AM) at a temperature of 21°C and allowed free access to water and food. The mice were individually acclimated to the metabolic chamber cage before measuring energy expenditure for at least 5 days, and then data were collected for another 24 hours. For Glucose tolerance test (GTT), mice were fasted overnight (18 h) before intraperitoneal injection of D-glucose (2 g/kg body weight). Subsequently, the clearance of plasma glucose was monitored following glucose administration. For insulin tolerance test, mice were fasted for 4 h before intraperitoneal injection of insulin (0.75 U/kg body weight). Every glucose was examined with tail-vein blood at indicated intervals (15, 30, 60, 90 and 120 min) after injection using a glucometer. For analyzing metabolic parameters, insulin (90080, Crystal Chem, Elk Grove Village, IL) were determined. For body composition measurement, fat and lean mass of each mouse in this study were measured by an EchoMRI100V, quantitative nuclear resonance system (Echo Medical Systems, Houston, TX).

#### Designer Receptors Exclusively Activated by Designer Drugs (DREADD) experiments

For DREADD experiments, 6 to 7 weeks old mice were injected either AAV-hSyn-DIO-mCherry or AAV-hSyn-DIO-hM4Di-mCherry with AAV-mGabra5-EGFP-Cre (3.8 x 10^13^ GC/ml) into the lateral hypothalamus of C57BL/6J. Following a recovery period of 3 weeks, Mice were given 5mg/kg/day CNO in drinking and high-fat diet food (TD 06414, Envigo) for 5 weeks in order to measure body weight and food consumption every week. CNO was dissolved in distilled water and protected from the light.

#### Quantitative real-time RT-PCR

Quantitative real-time RT-PCR was carried out using SYBR Green PCR Master Mix as described previously (Kwak et al., 2020). Briefly, reactions were performed in triplets in a total volume of 10μl containing 10pM primer, 4μl cDNA, and 5μl power SYBR Green PCR Master Mix (Applied Biosystems). The mRNA level of each gene was normalized to that of *18s* mRNA. Fold-change was calculated using the 2−ΔΔCT method. The following sequences of primers were used for real-time RT-PCR. The followings are the sequences of utilized primers.

*18S* forward: 5′- TGGCTC ATTAAATCAGTTATGGT -3′;

*18S* reverse: 5′- GTCGGCATGTATTAGCTCTAG -3′.

*Ucp1* forward: 5’- ACTGCCACACCTCCAGTCATT -3’;

*Ucp1* reverse: 5’- CTTTGCCTCACTCAGGATTGG -3’.

*Cidea* forward: 5’- TTCAAGGCCGTGTTAAGGAATC -3’;

*Cidea* reverse: 5’- CCAGGAACTGTCCCGTCATC - 3’.

*Prdm16* forward: 5’- CAGCACGGTGAAGCCATTC -3’;

*Prdm16* reverse: 5’- GCGTGCATCCGCTTGTG -3’

*Pgc1a* forward: 5’- AACCACACCCACAGGATCAGA -3’;

*Pgc1a* reverse: 5’- TCTTCGCTTTATTGCTCCATGA - 3’

*Dio2* forward: 5’ - CCACCTGACCACCTTTCACT - 3’;

*Dio2* reverse: 5’- TGGTTCCGGTGCTTCTTAAC -3’

*Adrb3* forward: 5’- CGACATGTTCCTCCACAAATCA -3’;

*Adrb3* reverse: 5’- TGGATTCCTGCTCTCAAACTA ACC- 3’

#### Preparation of gene-specific shRNA and shRNA virus

The shRNA sequences for scrambled, MAOB were adopted from previous studies (Nam et al., 2020; Yoon *et al*., 2014). The shRNA sequence for Gabra5 was designed with BLOCK-iT RNAi Designer (Invitrogen, USA) and cloned into pSicoR lentiviral vectors as previously described (Woo et al., 2012). pSicoR vectors were utilized for plasmid-based shRNA expression *in vitro.* Gabra5 shRNA was prepared from Human embryonic kidney 293T (HEK293T) cell which were purchased from ATCC (#CRL-3216, ATCC). Cell were cultured in Dulbecco’s modified Eagle’s medium (DMEM, Gibco, USA) supplemented with 25 mM of glucose, 4 mM of L-glutamine, 1 mM of sodium pyruvate, 10% heat-inactivated fetal bovine serum (#10082-147, Gibco) and 10,000 units/ml penicillin-streptomycin (#15140-122, Gibco). Cultures were maintained at 37°C in a humidified atmosphere containing 95% air and 5% CO2. Cells were transfected with DNA clone by transfection reagent (Effectene, #301425, Qiagen). Every construct was verified with sequencing after cloning. Cloned shRNA constructs were packaged into Lenti viruses in IBS Virus Facility (http://ibs.re.kr/virusfacility) and KIST Virus Facility (http://virus.kist.re.kr).

Sequence of scrambled shRNA for control: 5’-TCGCATAGCGTATGCCGTT-3’

Antisense sequence of shRNA target for MAOB: 5’-AATCGTAAGATACGATTCTGG-3’

Antisense sequence of shRNA target for Gabra5: 5’-CTTAAACCGCAGCCTTTCATC-3’

#### PRV inoculation

Recombinant pseudorabies virus (PRV) inoculation was performed in a biosafety level-2 operating room. For polysynaptic and retrograde circuit mapping from interscapular BAT and inguinal WAT, mice were anesthetized with vaporized isoflurane and settled in a brain stereotaxic apparatus (RWD Life Science Co.). After the interscapular BAT or inguinal WAT were exposed, two 500-nl injections of a PRV-CAG-EGFP were made into the brown fat or white fat on one side using a Hamilton syringe (KD Scientific). Shortly after surgery, mice were translocated to their home cages. After 5 days of microinjection, the animals were deeply anesthetized and then transcardially perfused and processed for immunohistochemistry.

#### Drug administration

KDS2010, a MAOB inhibitor, was synthesized as previously descried (Park et al., 2019). To test the effect of KDS2010 in HFD mice, we administered KDS2010 for 7-8 weeks through dissolving the compound in drinking water. The amount of KDS2010 was calculated as 20 mg/kg daily. Selegiline was dissolved in drinking water as calculated at 10 mg/kg daily as previously described (Jo *et al*., 2014). To compare the effect of KDS1524 with KDS2010, mice were orally administered 30 mg/kg for each drug for 15 days.

### QUANTIFICATION AND STATISTICAL ANALYSIS

All analysis were done blindly. The number of experimental samples, mean and SEM values are listed in Table S1. The numbers and individual dots refer to the number of cells unless otherwise clarified in figure legends. For data presentation and statistical analysis, Graphpad Prism (GraphPad Software) was used. For electrophysiology analysis, Minianalysis (synaptosoft) and Clampfit (Molecular Devices) were used. For CLAMS analysis, Oxymax for Windows software was used. For image analysis, Imagej (NIH, ver. 1.52s.) and IMARIS (Version 9.0.1, Oxford Instruments) softwares were used. Statistical significance was set at *p < 0.05, **p < 0.01, ***p < 0.001, ****p < 0.0001.

---

**Figure S1.**
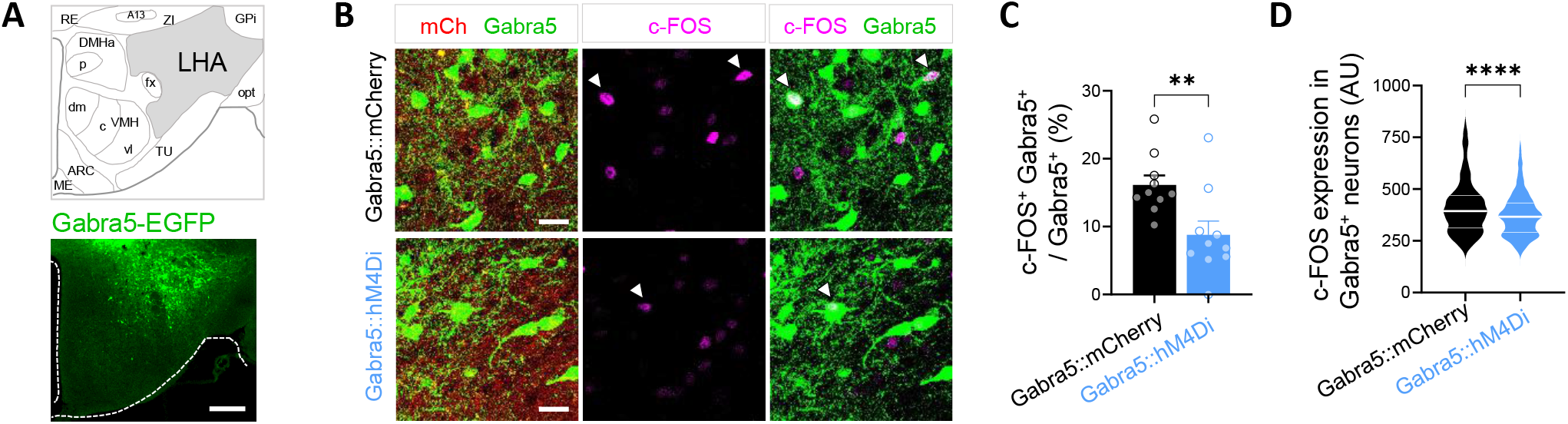
Inhibition of GABRA5^LHA^ neurons by DREADD. Related to Figure 2. (A) Representative images of AAV-Gabra5-EGFP-cre expression in LHA. Scale bar, 100 μm. (B) Gabra5:mCherry mice were injected with AAV-Gabra5-EGFP-cre and cre-activatable AAV carrying mCh (AAV-hSyn-DIO-mCh). Gabra5::hM4Di mice were injected with AAV-Gabra5-EGFP-cre and AAV-hSyn-DIO-hM4Di-mCh. Both groups were CNO drinking. c-FOS is shown as Magenta. Scale bar, 20 μm. (C and D) Percentage of c-FOS and Gabra5-double-positive neurons was lower in Gabra5::hM4Di than Gabra5::mCherry mice (n = 3-4 mice per group, n= 10, 10 slices, respectively). c-FOS expression in Gabra5-positive neuron was lower in Gabra5::hM4Di than Gabra5:mCherry mice. (n = 3-4 mice per group, n = 638, 464 cells, respectively).

**Figure S2.**
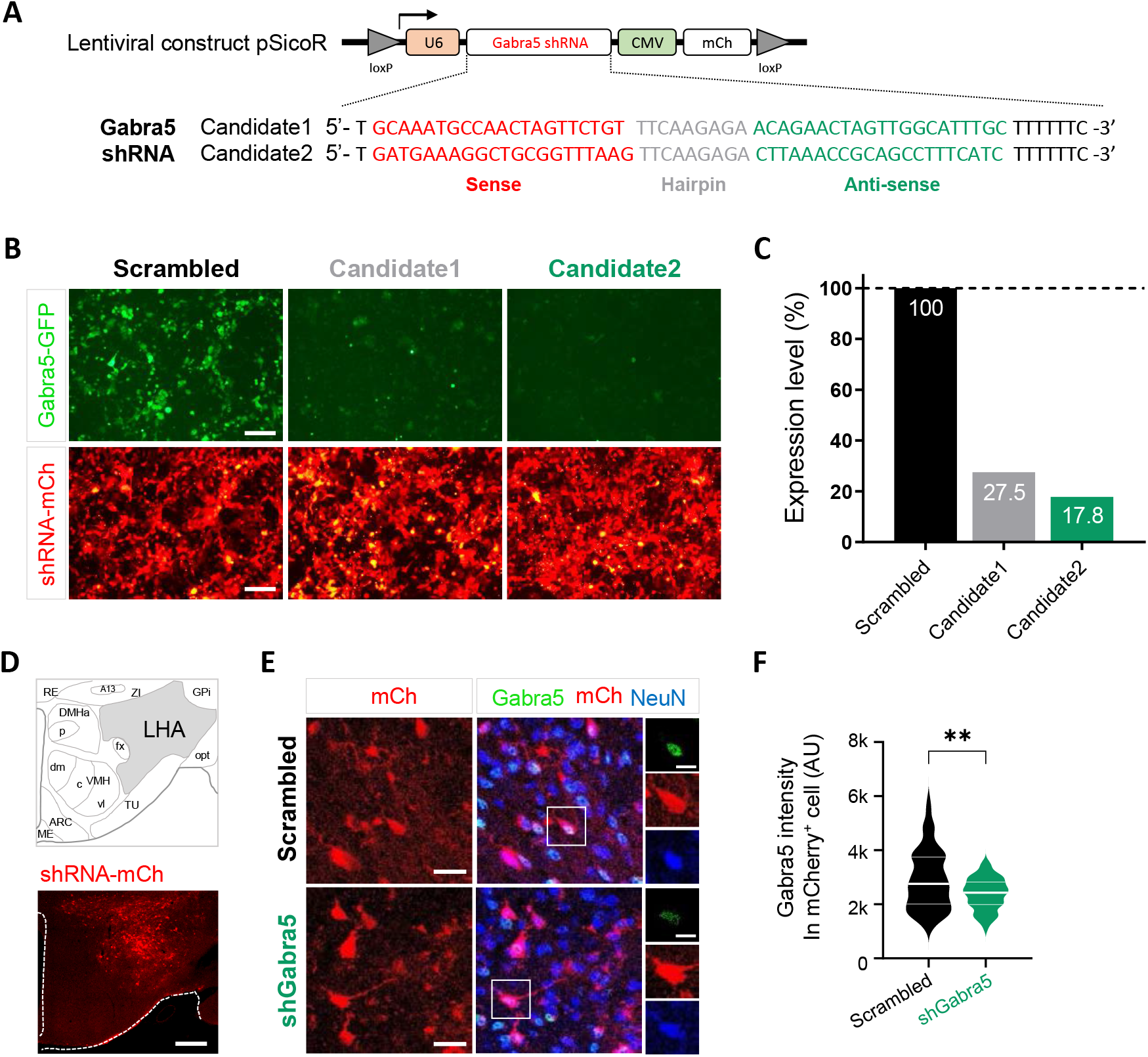
Development and validation of shRNAs for Gabra5 *in vitro.* and *in vivo.* Related to Figure 3. (A) Candidate sequences for Gabra5 shRNA are shown with vector information. Each shRNA was cloned in pSicoR vector to express under U6 promoter together with a cytomegalovirus (CMV) promoter driving expression of mCherry reporter gene. (B) Fluorescence images of HEK293-T cells co-transfected with Gabra5 shRNA (shRNA-mCh) with Gabra5 full clone (Gabra5-GFP). Top, Scrambled shRNA-mCherry was co-transfected with Gabra5-GFP. Bottom, Candidate 1 or candidate 2 of Gabra5 shRNA was co-transfected with Gabra5-EGFP. Scale bar, 100 μm. (C) Knock-down rate of Gabra5 shRNA candidates comprared to scrambled shRNA by RT-PCR. (D) Tom, Bottom, Gabra5-shRNA carrying lenti virus express mCherry in LHA. Scale bar, 100 μm. (E) Gabra5-IR was shown in GFP. Scale bar, 20 μm. Scale bar, 10 μm. (F) Expression of Gabra5 in mCherry-positive cell was significantly lower in shGabra5 mice compared to Scrambled mice (n= 96, 64 cells, respectively).

**Figure S3.**
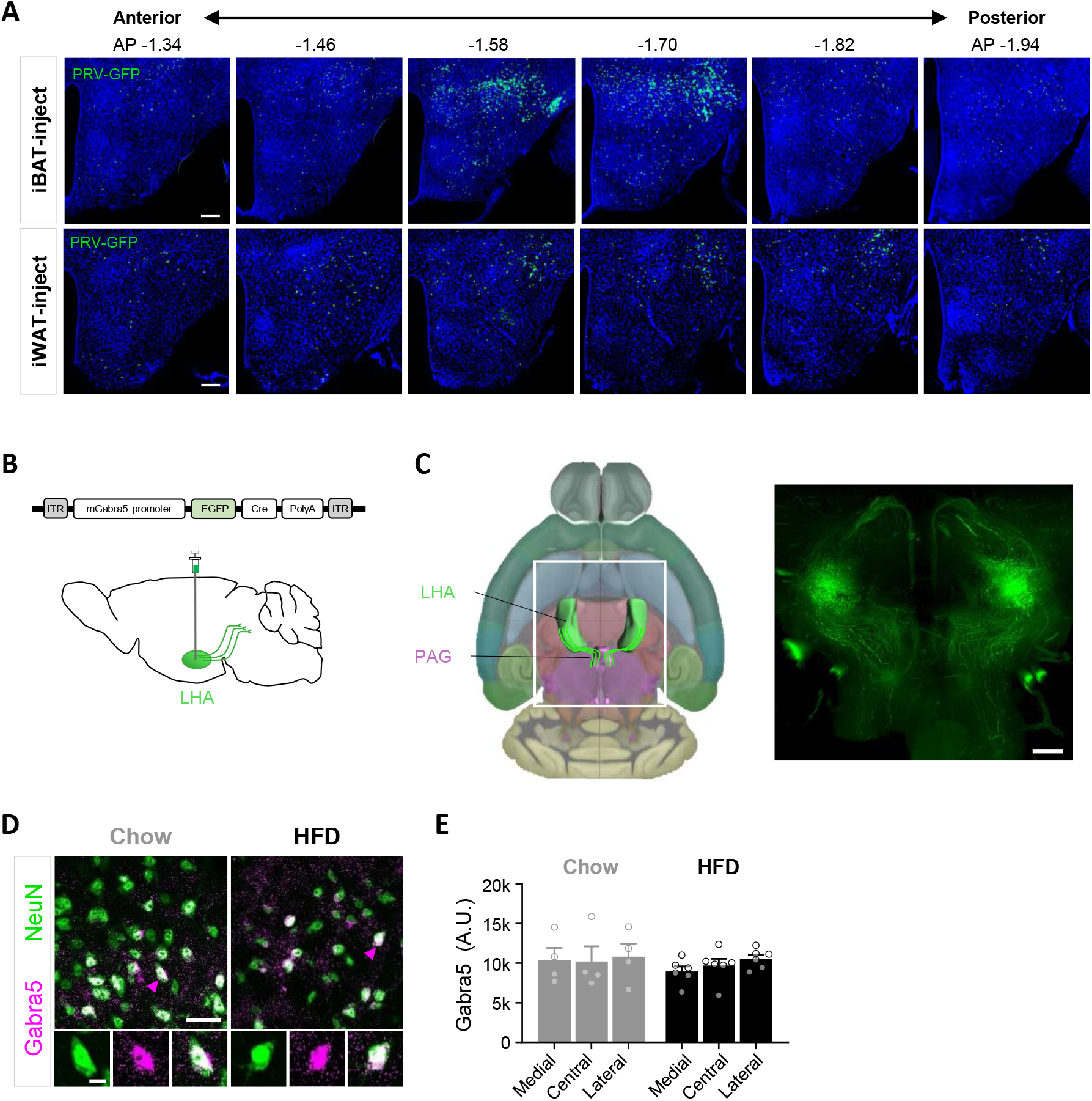
Expression of iBAT- and iWAT innervating neurons in LHA and expression of GABRA5^LHA^. Related to Figure 3, 4 and Supplemental Video 1. (A) We injected PRV-EGFP virus in iBAT and iWAT. After 5 days, mice showed robust expression of GFP-positive cells along the AP axis of LHA, from AP −1.34 mm to AP −1.94 mm. Scale bar, 100 μm. (B) Experimental schema of AAV-Gabra5-EGFP-cre virus injection and sagittal view of mouse brain. (C) Left, ventral view of the mouse brain from 3D Allen brain atlas. LHA is colored by green and PAG is colored by magenta. White rectangle indicates region of interest. Right, ventral image of LHA-injected mice with AAV-Gabra5-EGFP-cre virus. Scale bar, 300 μm. (D) Gabra5 expression in LHA of Chow and HFD mice. (E) The expression level of Gabra5 in medial, central and lateral part of the LHA was not significantly changed in HFD mice compared to Chow mice. Scale bar, 20 μm. Scale bar, 5 μm.

**Figure S4.**
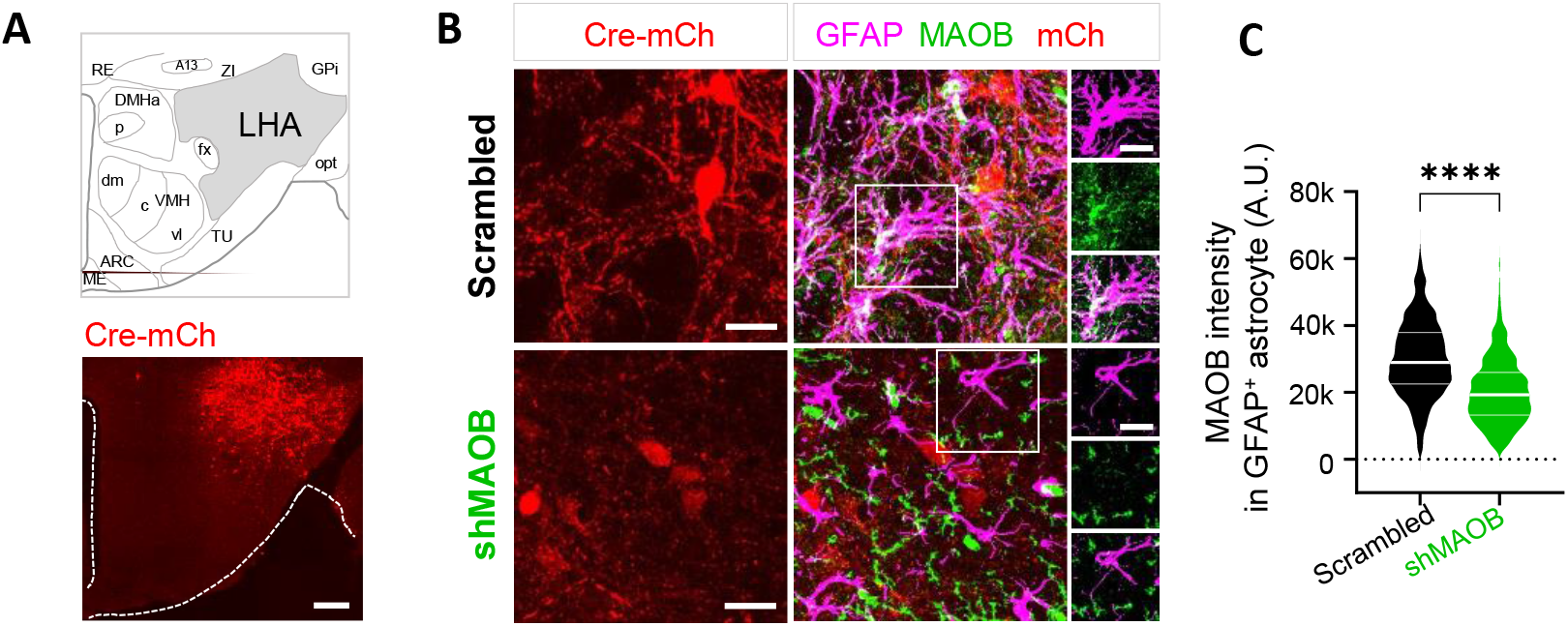
Knockdown of astrocytic GABA by gene-silencing. Related to Figure 5. (A) Top, schematic diagram. Bottom, Gabra5-shRNA carrying lenti virus express mCherry in LHA. Scale bar, 100 μm. (B) MAOB-IR was shown in GFP. Scale bar, 20 μm. Scale bar, 15 μm. (C) Expression of MAOB in GFAP-positive astrocyte was significantly lower in shMAOB mice compared to Scrambled mice (n=697, 345 cells, respectively).

**Figure S5.**
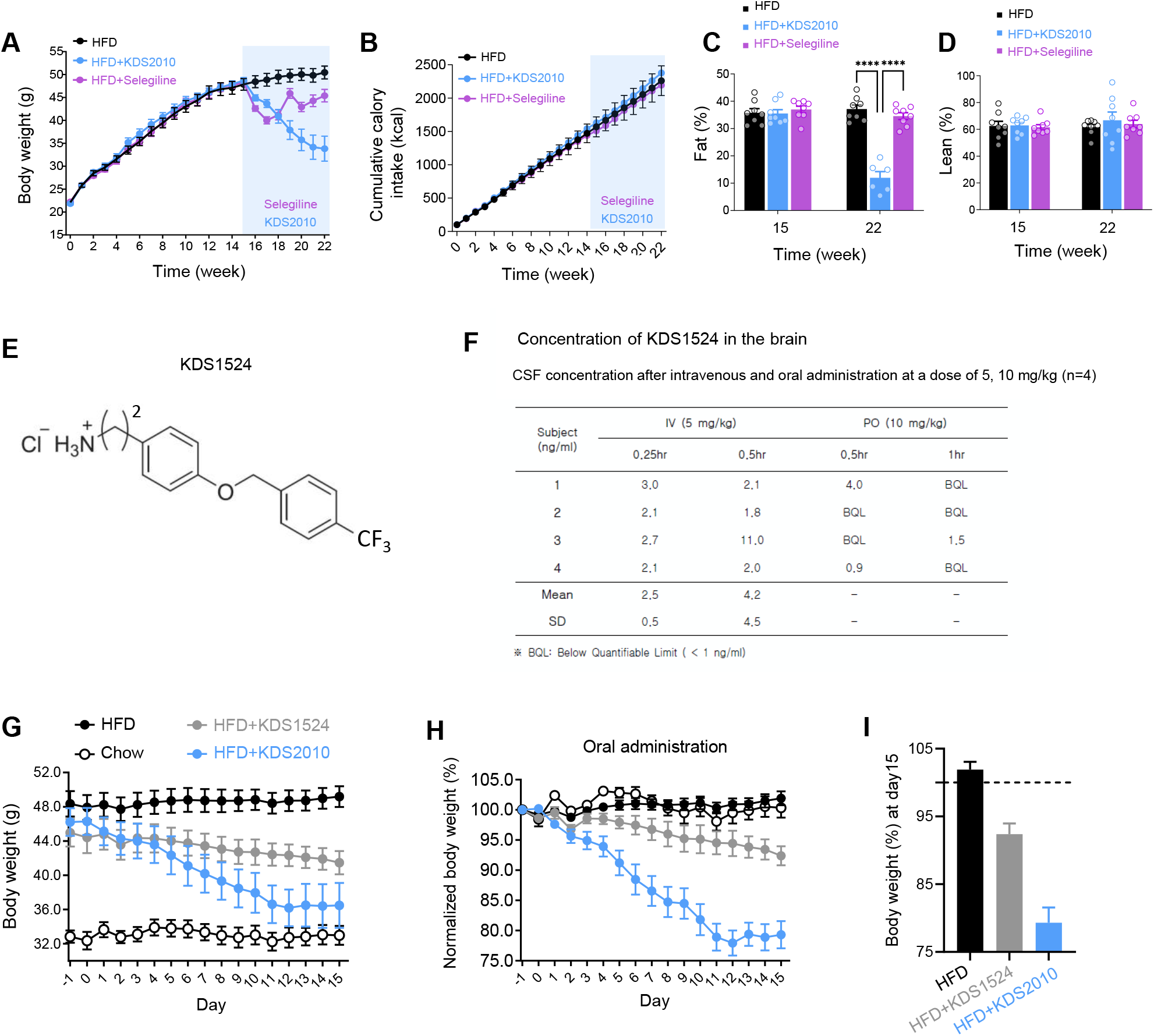
An irreversible MAOB inhibitor, Selegiline and a less-BBB-permeable MAOB inhibitor, KDS1524. Related to Figure 6. (A) Curves representing the kinetics of change in body weight among HFD, HFD with KDS2010, and HFD with selegiline mice over the 22 weeks following HFD treatment. Light blue box means KDS2010 or selegiline treatment in drinking water. n= 8 mice per group. (B) Cumulative food intake in HFD, HFD with KDS2010, HFD with selegiline mice over the 22 weeks. n=8 mice per group. (C and D) Quantification of percentage change of fat mass (C) and lean mass (D) at before (15 week) and after (22 week) KDS2010 or selegiline treatment. (E) Chemical structure of KDS1524. (F) Cerebrospinal fluid (CSF) concentration after intravenous and oral administration at a dose of 5, 10 mg/kg of KDS1524 (n= 4 mice). KDS1524 cannot pass the blood-brain-barrier. (G) Curves representing the kinetics of change in body weight in gram of chow, HFD, HFD with KDS1524 and HFD with KDS2010 mice over 16 days following HFD treatment. KDS2010 or KDS1524 was administered by oral injection (n= 8, 8, 8, 6 mice, respectively). (H) Curves representing the kinetics of change in body increase in percentage of chow, HFD, HFD with KDS1524 and HFD with KDS2010 mice. (I) Quantification of percentage change of body increase of HFD with KDS1524 and HFD with KDS2010 compared to HFD mice.

**Figure S6.**
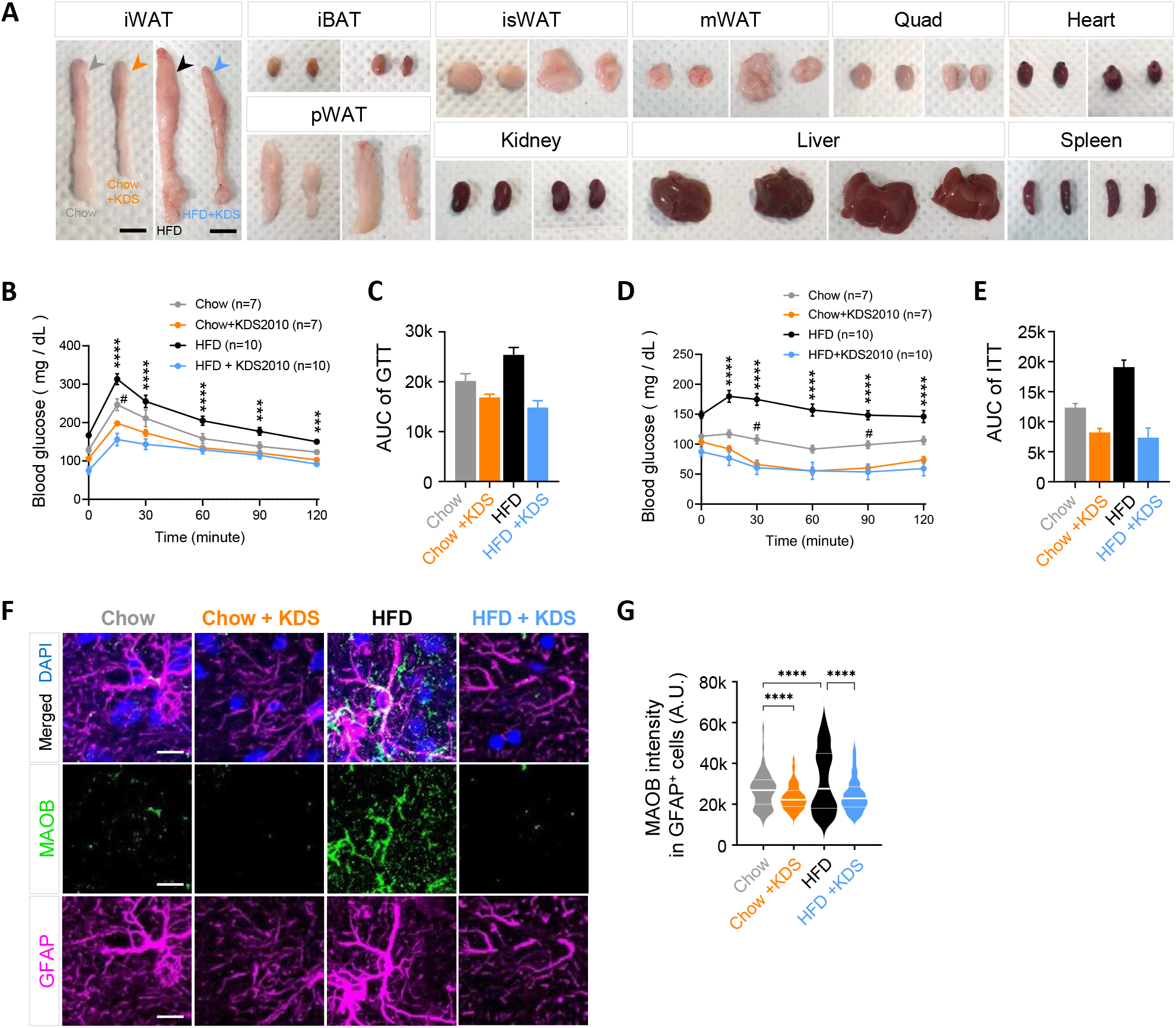
KDS2010 reduces fats, glucose tolerance and astrocytic GABA. Related to Figure 7. (A) Representative images of each organ of chow, chow with KDS2010, HFD, HFD with KDS2010 mice. Scale bar, 1 cm. (B) Time course of blood glucose levels during the oral glucose tolerance test (GTT) of chow, chow with KDS2010, HFD, HFD with KDS2010 mice. The mean glucose concentrations during the GTT were compared using Tukey’s multiple comparisons test at different time points (30, 60, 90 and 120 min). (***p<0.001,****p<0.0001, HFD vs. HFD with KDS2010; #p<0.05, Chow vs. Chow with KDS2010) n = 7-10 mice per group. (C) Quantification of area under curve (AUC) of glucose tolerance test. (D) Time course of blood glucose levels during the insulin tolerance test (ITT) of chow, chow with KDS2010, HFD, HFD with KDS2010 mice. The mean glucose concentrations during the ITT were compared using Tukey’s multiple comparisons test at different time points (30, 60, 90 and 120 min). (****p<0.0001, HFD vs. HFD with KDS2010; #p<0.05, Chow vs. Chow with KDS2010) n = 7-10 mice per group. (E) Quantification of area under curve (AUC) of insulin tolerance test. (F) Immunostaining for MAOB and GFAP in LHA of chow, chow with KDS2010, HFD and HFD with KDS2010 group. Scale bar, 10 μm. (G) MAOB intensity in GFAP-positive cells in chow, chow with KDS2010, HFD and HFD with KDS2010 group. n = 3-4 per group, n = 464, 463, 383, 296 cells, respectively.

**Figure S7.**
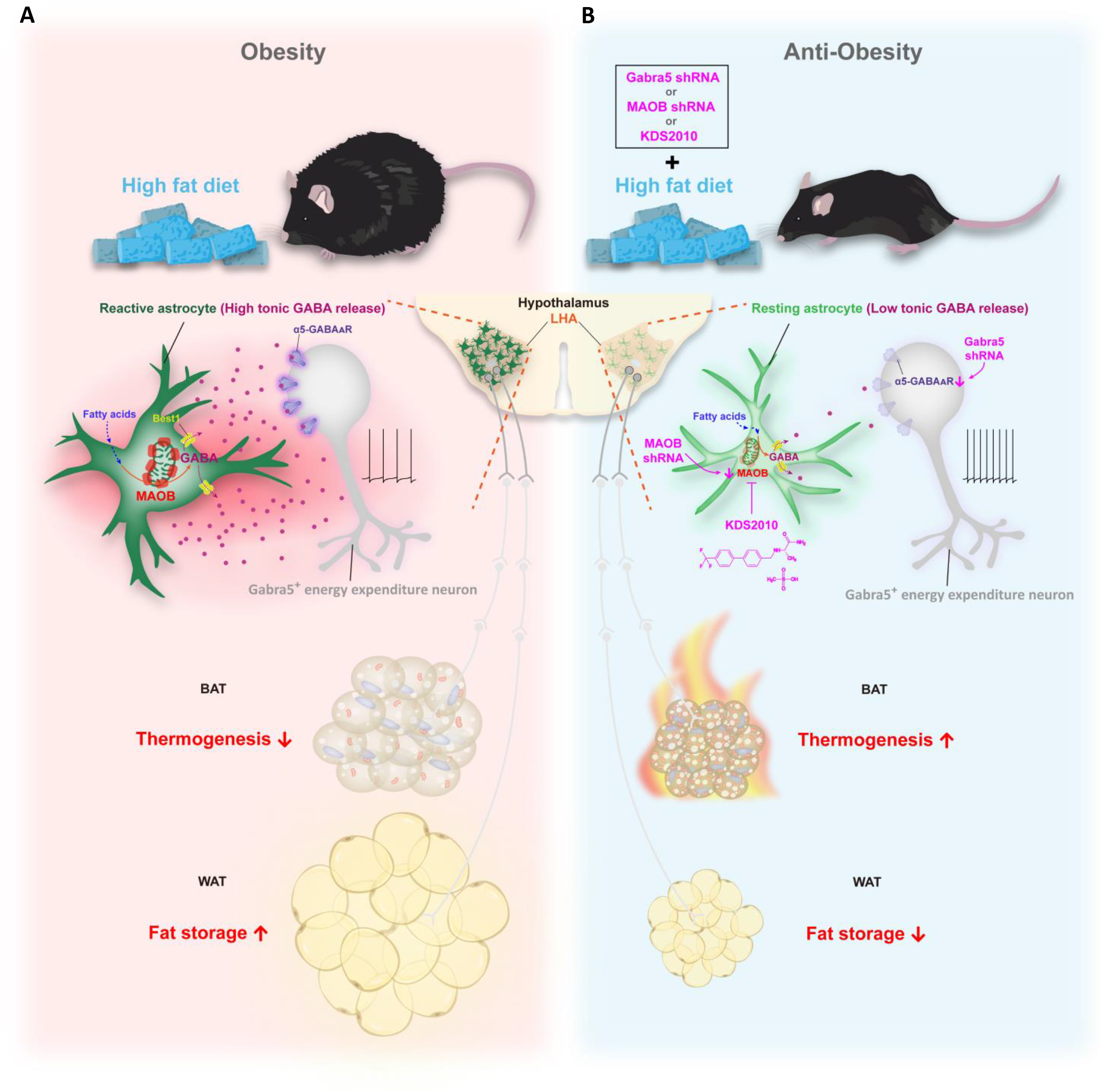
Summary. Related to Discussion part. (A) In DIO mouse model, GABA-synthesizing enzyme, MAOB, increased in reactive astrocytes of LHA. Chronic HFD leads to elevated MAOB to synthesize more GABA in reactive astrocytes, may tonically suppress the activity of GABRA5^LHA^. Inhibition of pacemaker firing GABAergic GABRA5^LHA^ attenuate energy expenditure and finally exacerbate obesity by decreasing thermogenesis and lipolysis-related gene expression in BAT and WAT. (B) Whereas the elevated tonic GABA or MAOB in hypothalamus of DIO mouse model, can be restored to the normal level by MAOB knockdown or Gabra5 knockdown or KDS2010 treatment. Emancipation of inhibition in pacemaker firing of GABRA5^LHA^ facilitates energy expenditure and then prevents obesity.

## Notes

### Competing Interest Statement

The authors have declared no competing interest.

### Summary of Updates

We changed the title and added one author.

